# Acetaminophen alters endogenous lipid signaling and attenuates pathological pain through a mechanism requiring diacylglycerol lipase, monoacylglycerol lipase and cannabinoid CB1 receptors in mice

**DOI:** 10.1101/2025.05.08.652937

**Authors:** Carlos Henrique Alves Jesus, Jonah L. Wirt, Taylor Woodward, Luana Assis Ferreira, Wenwen Du, John Hainline, Mirjam Huizenga, Lara Rems, Alex Makriyannis, Jenna A Himelstein, Noah M. Brauer, Mario van der Stelt, Heather B. Bradshaw, Andrea G. Hohmann

## Abstract

Acetaminophen (APAP) produces analgesia through mechanisms that remain poorly understood. Here, we tested the hypothesis that APAP-induced suppression of pathological pain is associated with cannabinoid receptors and activity of enzymes regulating endogenous lipids (diacylglycerol lipase, DAGL; monoacylglycerol lipase, MAGL) including endocannabinoids. APAP suppressed mechanical hypersensitivity in mouse models of post-surgical and inflammatory pain and the DAGL inhibitors (RHC-80267, DO34) and MAGL inhibitor JZL184 blocked APAP’s analgesic effects. Global (rimonabant, AM251) but not peripherally restricted (AM6545) cannabinoid receptor antagonists prevented APAP-induced analgesia. APAP increased corticosterone levels >2-fold and reduced prostaglandins >5-fold across the brain and in the paw skin. In addition, APAP reduced up to 39% of signaling lipids detected in the targeted screen in CFA-treated subjects in a tissue-dependent manner. These observations suggest that APAP plays a wider role in endogenous lipid signaling than previously hypothesized and provides novel insight into mechanisms of action.

## 1. Introduction

Acetaminophen (N-acetyl-para-aminophenol, APAP, or paracetamol) has been used as a pain and fever reliever for over a century. It is usually prescribed when contraindication for non-steroidal anti-inflammatory (NSAIDs) drugs exist [1]. Multiple targets contribute to *in vitro* and *in vivo* pharmacology of APAP, but its analgesic mechanism remains unclear. APAP poisoning is the most common pharmaceutical overdose worldwide, a trend largely driven by its widespread perception as a safe medication. In the United States, APAP toxicity remains the leading cause of acute liver failure [2–5]. Thus, a clearer understanding of its mechanisms of action could inform the development of more targeted analgesic therapies while reducing the risk of toxicity.

The endocannabinoid system contains multiple targets for drug development, including cannabinoid type 1 (CB1) and type 2 (CB2) receptors, endocannabinoids (e.g., anandamide (AEA) and 2-arachidonoyl glycerol (2-AG)) and biosynthetic (e.g., diacylglycerol lipase (DAGL)) and metabolic (e.g., fatty-acid amide hydrolase (FAAH) and monoacylglycerol lipase, (MAGL)) enzymes [6–14]. The endocannabinoid system has been implicated in the mechanism of action of APAP (for review see [15]) based largely upon studies evaluating acute nociceptive pain. Effects of APAP on a zymosan-induced inflammatory pain model were also eliminated in CB1 knock-out (KO) mice [16] and by CB1 antagonist treatment [17], suggesting APAP may activate CB1 via an indirect mechanism *in vivo*. Although APAP has been hypothesized to modulate DAGL [18], no direct or indirect evidence links this mechanism to the *in vivo* analgesic effects of APAP in models of pathological pain.

Most reports linking APAP to the endocannabinoid system have relied on *in vitro* systems or *in vivo* models of acute nociception (e.g., hot plate test), which lack strong translational relevance [19]. Here we tested that hypothesis that APAP-induced analgesia is associated with cannabinoid receptor signaling (e.g., CB1) and enzymes involved in the modulation (e.g., DAGL, MAGL, FAAH) of lipids such as 2-AG and AEA. To test this hypothesis, we used two mechanistically distinct preclinical mouse models of pathological pain: a hind paw incision model of post-operative pain and the complete Freund’s adjuvant (CFA) model of inflammatory pain to determine how APAP’s effects are modulated through enzyme and receptor modulation. Finally, we used high performance liquid chromatography tandem mass spectrometry LC MS/MS to evaluate modulation of a targeted small molecule lipidome (e.g., more than 100 endogenous lipids) by APAP in CFA-treated mice.

## 2. Methods

### 2.1 Subjects

Male and female mice on a C57BL/6 (Jackson Laboratories) or CD1 (CD1 IGS, also known as Crl:CD1(ICR)) background (Inotiv) were used. Mice were maintained in standard conditions of temperature (21 ± 2° C), humidity (45%) and under a light/dark cycle (12h/12h). All experimental procedures were approved by the Institutional Animal Care and Use Committee and followed the guidelines of the International Association for the Study of Pain. The experimenter was blinded to drug conditions in all studies.

### 2.2 Drugs and Chemicals

Acetaminophen (APAP, Sigma Aldrich, St. Louis MO, USA), RHC-80267 (Cayman Chemicals MI, USA) and JZL184 (Cayman) were purchased. DO34 and AM6545 were synthesized in the laboratories of Mario van der Stelt (Leiden University, Leiden, the Netherlands) and Alex Makriyannis (Center for Drug Discovery, Northeastern University, Boston, MA, USA), respectively.

RHC80267, DO34 and APAP were dissolved in a vehicle consisting of cremophor (Sigma-Aldrich, St. Louis, MO), ethanol (Sigma-Aldrich), and saline (Aquilite System, Hospira Inc, Lake Forest, IL) in a 1:1:18 ratio for intraperitoneal (i.p.) administration [13,20]. APAP was dissolved in a 1:1:8 ratio of dimethyl sulfoxide (DMSO), Kolliphore, and Mannitol for oral (p.o.) dosing. JZL-184 was dissolved in a 1:1:18 ratio of (DMSO), cremophor and saline [11,21]. Rimonabant was diluted in 20% DMSO and 1:1:8 ratio of ethanol, emulphor and saline [11]. AM251 was diluted in a solution of 1:1:1:17 ratio of DMSO, ethanol, Cremophor and saline [11]. AM6545 was dissolved in a 5: 2:2:16 ratio of DMSO, ethanol, emulphor and saline [9]. APAP was delivered (10 ml/kg) i.p. in the CFA model and via oral gavage in the incisional injury model. All other drugs were administered i.p. 30 minutes before APAP (p.o. or i.p.).

### 2.3 Incisional injury model

Male and female mice (CD1) were deeply anesthetized with isoflurane and a 5-mm longitudinal incision through the skin and fascia of the plantar region of the right paw [22]. After hemostasis with gentle pressure, the skin was closed with 8-0 nylon suture. Mechanical paw withdrawal thresholds were measured before (i.e., baseline) and one day after surgery to confirm incisional injury-induced mechanical hypersensitivity prior to assessments of drug efficacy.

### 2.4 CFA-induced inflammatory pain

Complete Freund’s adjuvant (CFA) (1:1 in saline)[23,24] was injected (20 µl) into the plantar surface (i.pl.) of the right hind paw of male (C57BL/6) or female (CD1) mice. Mechanical paw withdrawal thresholds were measured before (i.e., baseline) and two days after CFA injection to confirm presence of adjuvant-induced mechanical hypersensitivity prior to pharmacological manipulations.

### 2.5 Assessment of paw withdrawal thresholds

Paw withdrawal thresholds (in grams) were measured using an electronic von Frey aesthesiometer (IITC Life Science, Woodland Hills, CA) as described [11,23,24]. Mice were habituated (≥30 minutes) to inverted plexiglass chambers placed on a metal mesh. Withdrawal thresholds were recorded in duplicate for each hind paw with a 2-minute interval between stimulations to provide a single value per paw/per animal/per timepoint. Withdrawal thresholds were measured before and after injury and at various timepoints (i.e., 30, 90, 240 min) following pharmacological manipulations.

### 2.6 Assessment of peripheral edema

Thickness (in mm) of each hind paw was measured in duplicate using an electronic caliper (World Precision Instruments, INC; Sarasota, FL, USA; Digital Caliper 501601) as described [25]. Measurements were averaged for each paw before and after i.pl. and i.p. injections in CFA-injected mice.

### 2.7 Locomotor activity

CD1 mice were habituated to the test room in their home cages (≥1 hour) before being placed in an Omnitech Superflex Node activity meter (42 × 42 x 30 cm). Total distance travelled (cm) during a 30 min testing period was calculated using Fusion 6.5 software (Omnitech Electronics, Columbus, OH). Mice were habituated to the activity meter (30 min) 24 h prior to pharmacological manipulations. Locomotor activity was assessed for 30 minutes after drug treatments (i.p.). Time in the center of the apparatus (20 x 20 cm), rest time (inactivity ≥1 sec), vertical time (time breaking beams in the vertical layer) and total distance traveled were measured [10].

### 2.8 Body temperature

Body temperature (°C) was assessed using a thermometer (Physitemp Instruments, Inc., Clifton, NJ) and rectal probe (Braintree Laboratories, Inc., Braintree, MA) before (i.e., baseline) and after pharmacological treatments [7,10].

### 2.9 Tail-immersion test

The hot water tail immersion test was used to evaluate acute nociception. The distal portion (2 cm) of the mouse tail was submerged in a water bath (52°C) and the latency to tail flick was measured [7,10,26]. Three baseline values were obtained at 10-min intervals. Tail-flick latency was measured before and after drug treatments using a cut-off latency of 15 seconds.

### 2.10 Experimental procedures

*Experiment 1: Dose response of APAP in an incisional injury model.* We evaluated impact of APAP (100 and 300 mg/kg, p.o.) and vehicle on mechanical hypersensitivity induced by plantar paw incision using male mice.

*Experiment 2: Effect of DAGL and MAGL inhibition on anti-allodynic efficacy of APAP following incisional injury.* Separate cohorts of male and female mice with established incisional injury were pretreated with RHC-80267 (20 mg/kg), DO34 (30 mg/kg) or vehicle (i.p.) followed by APAP (300 mg/kg) or vehicle p.o. treatment. In a separate study, male mice with incisional injury received (i.p.) JZL-184 (4 mg/kg) or vehicle prior to APAP (300 mg/kg) or vehicle (p.o.) injection.

*Experiment 3: Effect of CB1 receptor antagonist on anti-allodynic efficacy of APAP following incisional injury.* Male and female mice with incisional injury received (i.p.) AM251 (5 mg/kg) or vehicle before APAP (300 mg/kg) or vehicle (p.o.) treatment.

*Experiment 4: Effect of APAP on body temperature, tail-flick latency and locomotor activity.* Tail-flick antinociception and rectal temperature were measured before (baseline) and after APAP (100 or 300 mg/kg) or vehicle (i.p.) treatment in otherwise naïve male mice. We also examined effects of pretreatment (i.p.) with AM251 (5 mg/kg), RHC-80267 (20 mg/kg), or their respective vehicles on responses induced by APAP in each assay. After a 7-day washout period, baseline activity meter assessments were performed. Then, 24 hours later, locomotor activity was reassessed after treatment (i.p.) with the same drug combination: Veh-APAP (300 mg/kg), RHC-80267 (20 mg/kg)-APAP, RHC-809267 (20 mg/kg)-Veh, and Veh-Veh. In a separate study, groups received (i.p.): Veh-APAP (300 mg/kg), AM251 (5 mg/kg)-APAP (300 mg/kg), AM251 (5 mg/kg)-Veh or Veh-Veh.

*Experiment 5: Dose response of APAP in CFA-induced inflammatory pain.* We evaluated impact of APAP (30, 100 and 300 mg/kg, i.p.) on mechanical hypersensitivity in male and female mice 2 days following unilateral intraplantar (i.pl.) injection of CFA.

*Experiment 6: Effect of DAGL and MAGL inhibition on the anti-allodynic effect of APAP in CFA-induced inflammatory pain.* We assessed the effect of DAGL inhibitors (RHC-80267 (20 mg/kg) and DO34 (30 mg/kg) and MAGL inhibitor JZL-184 (1.6 mg/kg), administered i.p., on APAP (300 mg/kg, i.p.)-induced analgesia in the CFA model in male and female mice.

*Experiment 7: Effect of CB1 receptor antagonists on the anti-allodynic effect of APAP in CFA model.* We examined the effect of pretreatment with global CB1 receptor antagonists (AM251 or Rimonabant, 5 mg/kg; i.p.) or vehicle on anti-allodynic effects of APAP (300 mg/kg, i.p.) in male and/or female mice injected unilaterally with CFA.

*Experiment 8: Effect of peripherally-restricted CB1 receptor antagonist on anti-allodynic effect of APAP in CFA model*. We assessed the effect of a peripherally-restricted CB1 receptor antagonist AM6545 (10 mg/kg; i.p.) or vehicle on the antinociceptive efficacy of APAP (300 mg/kg, i.p.) in male and female CFA-injected mice.

*Experiment 9: Effect of APAP on lipid levels in CNS.* We evaluated impact of APAP (300 mg/kg, i.p.) or vehicle on lipid levels in male mice injected (i.pl.) with CFA. Mice were sacrificed 90 minutes following injection (i.e., peak analgesic effect).

### 2.11 Tissue collection

Mice were decapitated without anesthesia and brain, spinal cord and paw skin were rapidly extracted and flash-frozen in liquid nitrogen [27]. Coronal brain sections were prepared using a pre-cooled stainless steel brain matrix, and tissue punches (1- or 2-mm diameter) were obtained from the periaqueductal gray (PAG), rostral ventromedial medulla (RVM), and thalamus (THA). Lumbar spinal cord (SC) segments were collected from the same mice. All samples were immediately weighed and stored (-80°C ) until further processing.

### 2.12 Lipid Extraction and High-Pressure Liquid Chromatography Coupled to Tandem Mass Spectrometry (HPLC/MS)

Lipid extraction and partial purification were performed on brain tissue punches as described previously [28,29]. Brain punches were added to 2 mL HPLC-grade methanol (MeOH) spiked with 10 uL of 1 uM d8-AEA (Cayman Chemical, Ann Arbor, MI, USA) and incubated on ice for 2 hours in the dark. After incubation, samples were sonicated for ∼30 seconds and centrifuged (19,000 G, 20 min, 20°C). Organic supernatants were added to 8 mL HPLC-grade water to make a 20% organic solution, which was partially purified using 500mg Bond Elut C18 solid phase extraction columns (Agilent Technologies, Santa Clara, CA, USA). Lipids were eluted with 1.5 mL of 65%, 75% an 100% methanol, which were stored at -80°C until MS analysis. From these methanolic extracts 106 lipids were screened using a SCIEX 7500 (SCIEX, Framingham, MA) coupled to a Shimadzu LC system LC-40DX3. Analytes were separated by a Luna 3 µm C18(2) 50 x 2 mm analytical column (Phenomenex, Torrence, CA) and were detected with a multiple reaction monitoring (MRM) method previously optimized for each analyte. Analyte concentrations were quantified with SCIEX OS peak matching software using standard curves from purchased standards (Cayman Chemical, Ann Arbor, MI) or those made-in house as described previously [30]. An MRM method for APAP and AM404 was created with standards purchased from Cayman Chemical. APAP and AM404 were detected in positive mode with a parent/daughter transition of 152.11/110.02 (APAP) and 396.33/152.02 (AM404). APAP was quantified by diluting the 65% percent methanolic extract obtained during solid phase extraction 100-fold in 20% methanol, and AM404 was screened in the undiluted 100% methanolic sample extracts. Peak areas in samples (if detected) were compared to a 6-point standard curve.

### 2.13 Statistical analysis

Behavioral data were analyzed by Two-way repeated measures or One-way ANOVA and Bonferroni’s *post hoc* test using GraphPad Prism 10 (GraphPad Software, La Jolla, CA). All statistical and descriptive data are available as Supplementary information (Supplemental Table 1). Exploratory lipidomic data was analyzed using a two-tailed t-test for each analyte in each tissue using a custom Python (version 3.12.4) script that also calculated fold change and Cohen’s d for each comparison. A summary heatmap was also generated using this code, with statistical significance (p<0.05) and trending comparisons (p values between 0.1 and 0.05) differentiated by color darkness as previously described [28–30].

## 3. Results

### 3.1 General experimental results

In all studies, incisional injury or intraplantar CFA reduced paw withdrawal thresholds relative to baseline (pre-injury). In general, withdrawal thresholds in the injured paw changed in a treatment and time-dependent manner and varied across time irrespective of drug treatment (**Supplemental Table 1**). None of the treatments reliably altered paw withdrawal thresholds in the non-injured (contralateral) paw (**Supplemental Table 1, Supplemental Figures 1A-B, 2A-D, 4A-H, 5A-F**).

### 3.2 APAP attenuated mechanical allodynia in a model of incisional injury

In male mice, APAP (100 and 300 mg/kg, p.o.) increased paw withdrawal thresholds overall and in a time-dependent manner in the injured (ipsilateral) paw (**Figure 1A**: Treatment: F _2,15_=8.477, p=0.0034; Interaction: F_6,45_=5.831, p=0.0001); both high (p< 0.0001) and low (p=0.0005) dose APAP increased paw withdrawal thresholds relative to Vehicle treatment (p<0.0001) but did not differ from each other overall (p=0.4698). No alterations in paw withdrawal thresholds were detected in the contralateral paw (**Figure 1B**).

**Figure 1.**
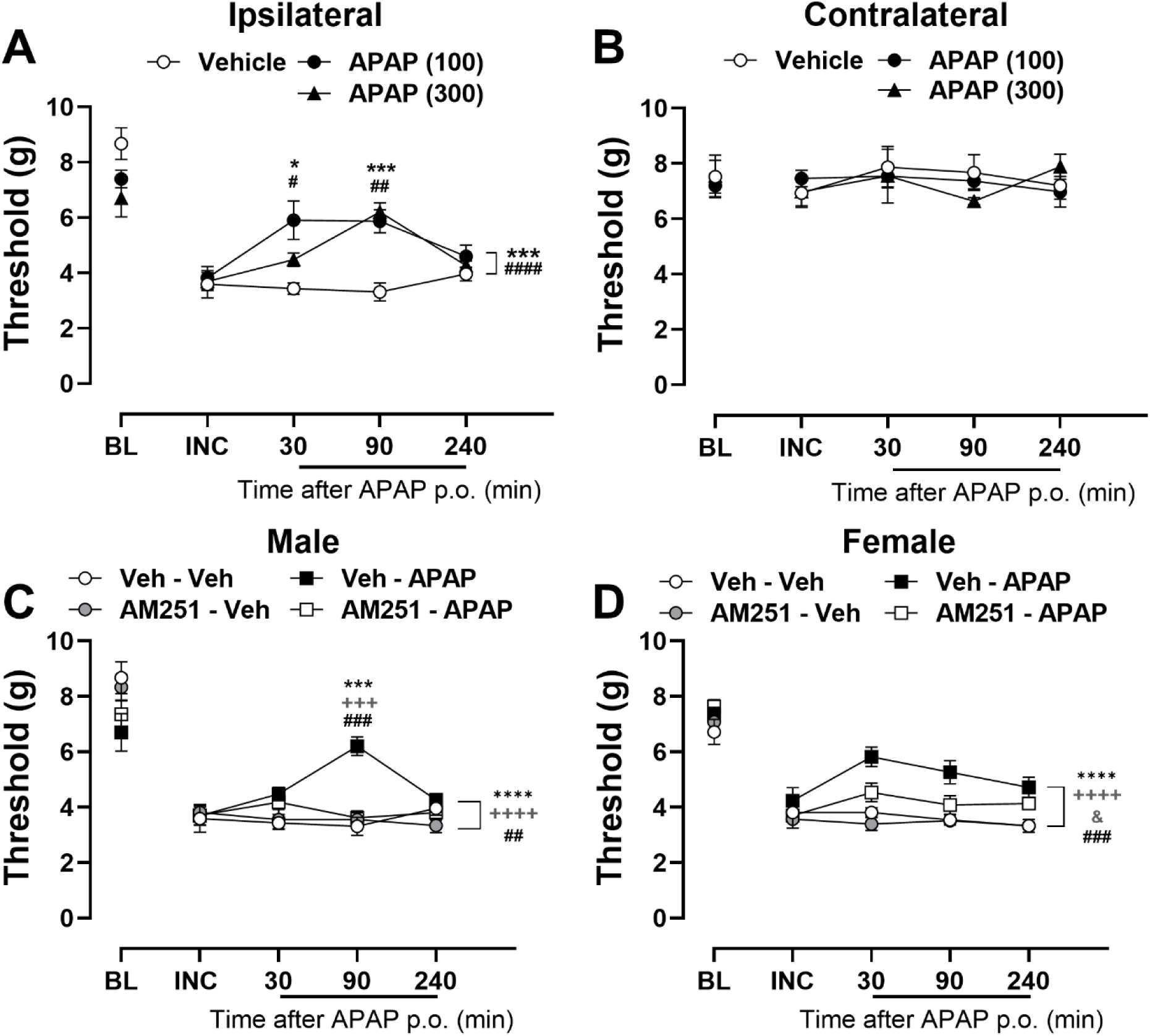
APAP suppresses mechanical hypersensitivity induced by incisional injury in the hind paw, an effect blocked by CB1 cannabinoid receptor antagonist AM251. APAP elevated mechanical paw withdrawal threshold in the (**A**) ipsilateral paw (injured paw) at all doses (100 and 300 mg/kg, p.o.) without altering responsiveness in the (**B**) contralateral paw in male mice. CB1 receptor antagonist AM251 (5 mg/kg, i.p.) suppresses the anti-allodynic effect of APAP (300 mg/kg, p.o.) evaluated in the ipsilateral paw of male (**C**) and female mice (**D**) with incisional injury (n=6 per group). Data show mean (± SEM) for ipsilateral and contralateral paws. ***Panel A-B****: Two-way ANOVA followed by Bonferroni’s post hoc test. Interaction between time points and treatment effect: * P <0.05, *** P <0.001 Vehicle vs. APAP 300; # P <0.05, ## P <0.01 Vehicle vs. APAP 100. Brackets (main treatment effect only): *** P <0.001 Vehicle vs. APAP 300; #### P <0.0001 Vehicle vs. APAP 100. **Panels C-D**: Two-way ANOVA followed by Bonferroni’s post hoc test. Interaction between time points and treatment effect: *** P <0.001 Veh-Veh vs. Veh-APAP; ### P <0.001, Veh-APAP vs. AM251-APAP; +++ P<0.001 AM251-Veh vs. Veh-APAP. Brackets (main treatment effect only): **** P <0.0001 Veh-Veh vs. Veh-APAP; ## P <0.01, ### P <0.001 Veh-APAP vs. AM251-APAP; ++++ P<0.0001 AM251-Veh vs. Veh-APAP; & p<0.05 AM251-Veh vs. AM251-APAP*.

### 3.3 Global CB1 receptor antagonist attenuated the anti-allodynic effect of APAP

In male mice with incisional injury, the CB1 receptor antagonist AM251 prevented the anti-allodynic effect of orally-administered APAP (**Figure 1C**; Treatment: F_3,20_=9.131, p=0.0005; Interaction: F_9,60_=4.293, p=0.0002). Veh-APAP (300 mg/kg p.o.) elevated withdrawal thresholds in the injured paw, relative to groups receiving Veh-Veh (p<0.0001), AM251-Veh (p<0.0001) or AM251-APAP (p=0.0020). Effects of AM251-APAP and AM251-Veh did not differ from Veh-Veh treatment (p>0.9999 for each comparison).

In female mice with incisional injury, AM251 similarly prevented the anti-allodynic effect of APAP (**Figure 1D**; Treatment: F_3,20_=35.91, p<0.0001; Interaction: F_9,60_=1.208, p=0.3074); Veh-APAP (300 mg/kg, p.o.) elevated paw withdrawal thresholds compared to Veh-Veh (p<0.0001), AM251-Veh (p<0.0001) or AM251-APAP (p=0.0005) whereas AM251-Veh (p>0.9999) and AM251-APAP (p=0.1583) did not differ from Veh-Veh treatment.

### 3.4 RHC-80267, DO34 and JZL 184 attenuate the anti-allodynic effect of APAP in a model of incisional injury

In male mice with incisional injury, the DAGL inhibitor RHC-80267 blocked the anti-allodynic effect of orally-administered APAP (**Figure 2A**; Treatment: F_3,20_=5.202, p=0.0081; Interaction: F_9,60_=3.210, p=0.0031); APAP (300 mg/kg p.o.) increased withdrawal thresholds in the ipsilateral (injured) paw overall relative to Veh-Veh (p<0.0001) or RHC-Veh (p=0.0051) and RHC80267 reliably blocked the anti-allodynic effect of APAP (p=0.0082). Effects of RHC-80267-APAP did not differ from Veh-Veh or RHC-80267-Veh treatments and did not differ from each other (p>0.9999 for each comparison).

**Figure 2.**
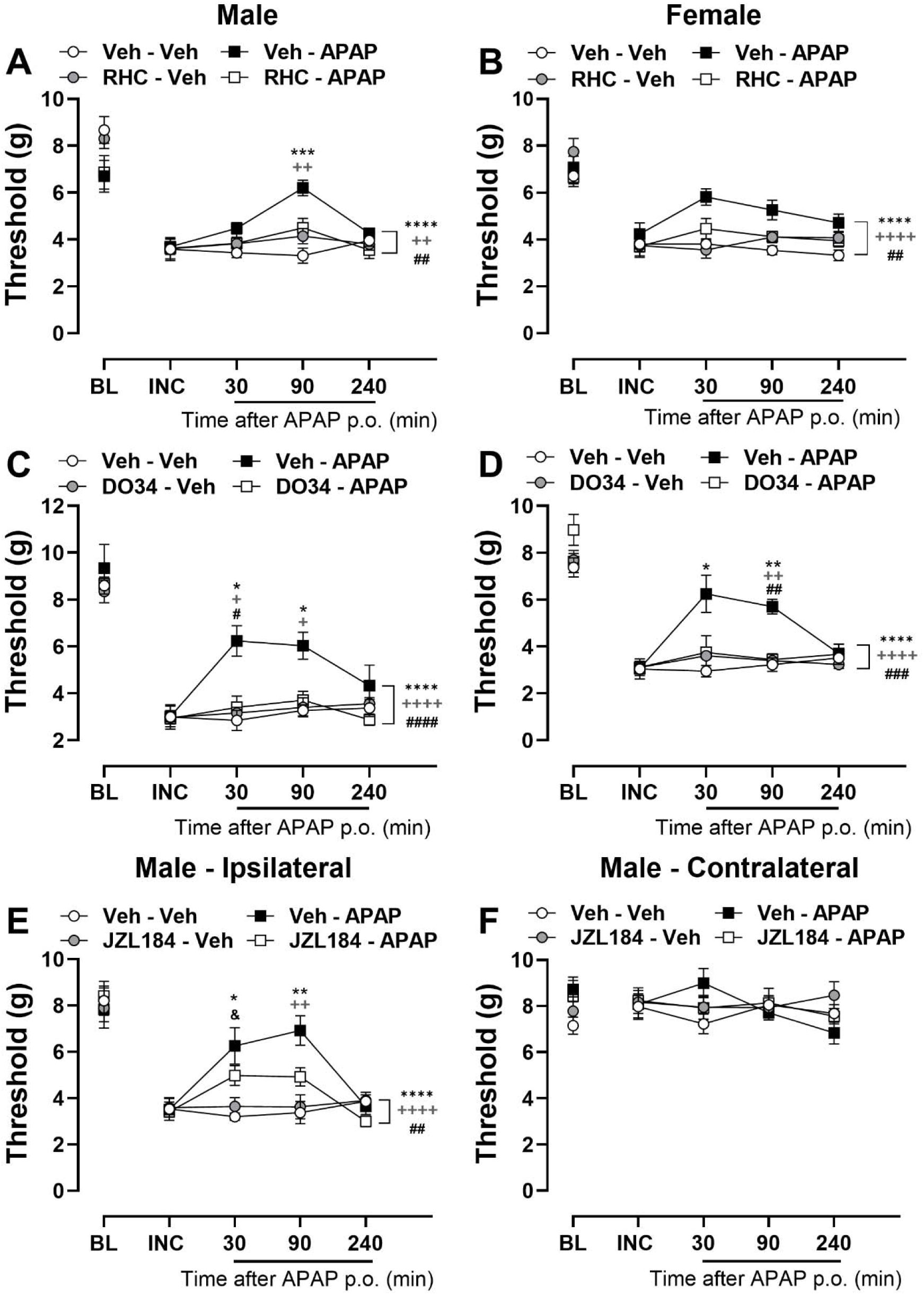
Anti-allodynic effects of APAP are suppressed by pharmacological inhibitors of DAGL and MAGL in a mouse model of post-surgical pain. DAGL inhibitor RHC-80267 (20 mg/kg, i.p.) suppresses the anti-allodynic effect of APAP (300 mg/kg, p.o) in the ipsilateral paw of (**A**) male and (**B**) female mice with incisional injury (n=6 per group). DAGL inhibitor DO34 (30 mg/kg, i.p.) suppresses the anti-allodynic effect of APAP (300 mg/kg, p.o) in the ipsilateral paw of (**C**) male and (**D**) female mice with incisional injury. MAGL inhibitor JZL 184 (4 mg/kg, i.p.) suppresses the anti-allodynic effect of APAP (300 mg/kg, p.o) in the ipsilateral paw (**E**) but not the contralateral paw (**F**) of male mice with incisional injury. Data show mean (± SEM) (n=6-8 per group). Two-way ANOVA followed by Bonferroni’s post hoc test. Interaction between time points and treatment effect: *\*P<0.05, **P <0.01, ***P <0.001 Veh-Veh vs. Veh-APAP; #P <0.05, ## P <0.01 Veh-APAP vs. RHC-APAP or Veh-APAP vs. DO34-APAP; + P<0.05, ++P<0.01 RHC-Veh vs. Veh-APAP or DO34-Veh vs. Veh-APAP or JZL 184-Veh vs. Veh-APAP. Brackets (main treatment effect only): **** P <0.0001 Veh-Veh vs. Veh-APAP; ## P <0.01, ### P <0.001, #### P <0.0001 Veh-APAP vs. RHC-APAP or Veh-APAP vs. DO34-APAP or Veh-APAP vs. JZL 184-APAP; ++ P<0.01, ++++ P<0.0001 RHC-Veh vs. Veh-APAP or DO34-Veh vs. Veh-APAP or JZL 184-Veh vs. Veh-APAP*.

In female mice with incisional injury, RHC-80267 blocked the anti-allodynic effect of orally-administered APAP (**Figure 2B**; Treatment: F_3,20_=10.07, p=0.0003; Interaction: F_9,60_=1.421, p=0.1995); Veh-APAP (300 mg/kg, p.o.) increased paw withdrawal thresholds in the injured paw compared to mice receiving Veh-Veh (p<0.0001) or RHC-80267-Veh (p<0.0001). RHC-80267 pretreatment, at a dose that did not differ from vehicle alone (p>0.9999), blocked the anti-allodynic effect of APAP (p=0.0011).

In male mice with incisional injury, the DAGL inhibitor DO34 blocked the anti-allodynic effect of APAP (**Figure 2C**; Treatment: F_3,20_=8.708, p=0.0007; Interaction: F_9,60_=3.387, p=0.0020); Veh-APAP (300 mg/kg, p.o.) increased withdrawal thresholds overall, in comparison to Veh-Veh (p<0.0001), DO34 (30 mg/kg)-Veh (p<0.0001) and DO34-APAP (p<0.0001) treatment. DO34-Veh did not differ from Veh-Veh or DO34-APAP (p>0.9999 for each comparison).

In female mice with incisional injury, DO34 (30 mg/kg) pre-treatment prevented the anti-allodynic effect of APAP (300 mg/kg, p.o.) (**Figure 2D**; Treatment: F_3,20_=8.442, p=0.0008;

Interaction: F_9,60_=4.388, p=0.0002); Veh-APAP elevated withdrawal thresholds in the injured paw compared to treatment with either Veh-Veh or DO34-Veh (p<0.0001 for each comparison) and pre-treatment with DO34 attenuated the anti-allodynic effect of APAP (p=0.0003). DO34-Veh did not differ from Veh-Veh or DO34-APAP (p>0.9999 for each comparison).

In male mice with incisional injury, a low dose of the MAGL inhibitor JZL184 attenuated the anti-allodynic effect of orally-administered APAP (**Figure 2E**; Treatment: F_3,28_=9.213, p=0.0002; Interaction: F_7.584,70.79_=5.151, p<0.0001). Veh-APAP (300 mg/kg, p.o.) increased paw withdrawal thresholds compared to Veh-Veh or JZL 184 (1.6 mg/kg, i.p.)-Veh (p<0.0001 for each comparison) and pre-treatment with JZL184 reduced the anti-allodynic effect of APAP (p=0.0079). Ipsilateral paw withdrawal thresholds in JZL184-Veh (p>0.9999) and JZL184-APAP (p>0.9999) groups were similar to Veh-Veh treatment. No differences in paw withdrawal thresholds were detected in the contralateral paw (**Figure 2F**).

### 3.5 CB1 receptor antagonist (AM251) does not attenuate side effects of high dose of APAP treatment in naïve mice

APAP-induced reductions in body temperature were not mediated by CB1 receptors (**Figure 3A**, Treatment: F_3,28_=18.37, p<0.0001; Interaction: F_9,84_=11.23, p<0.0001). APAP (300 mg/kg, i.p.) decreased body temperature compared with groups treated with vehicle or AM251 alone irrespective of AM251 pretreatment (p<0.0001 for all comparisons), which did not itself alter tail-flick latencies relative to vehicle (p=0.8661). None of the pharmacological treatments altered tail-flick latencies in otherwise naïve mice (**Figure 3B**). APAP did not reliably reduce total distance traveled irrespective of AM51 pretreatment and the interaction between time and treatment was not significant (**Figure 3C**, Treatment: F_3,27_=2.687, p=0.0664; Interaction: F_3,27_=0.7625, p=0.5250).

**Figure 3.**
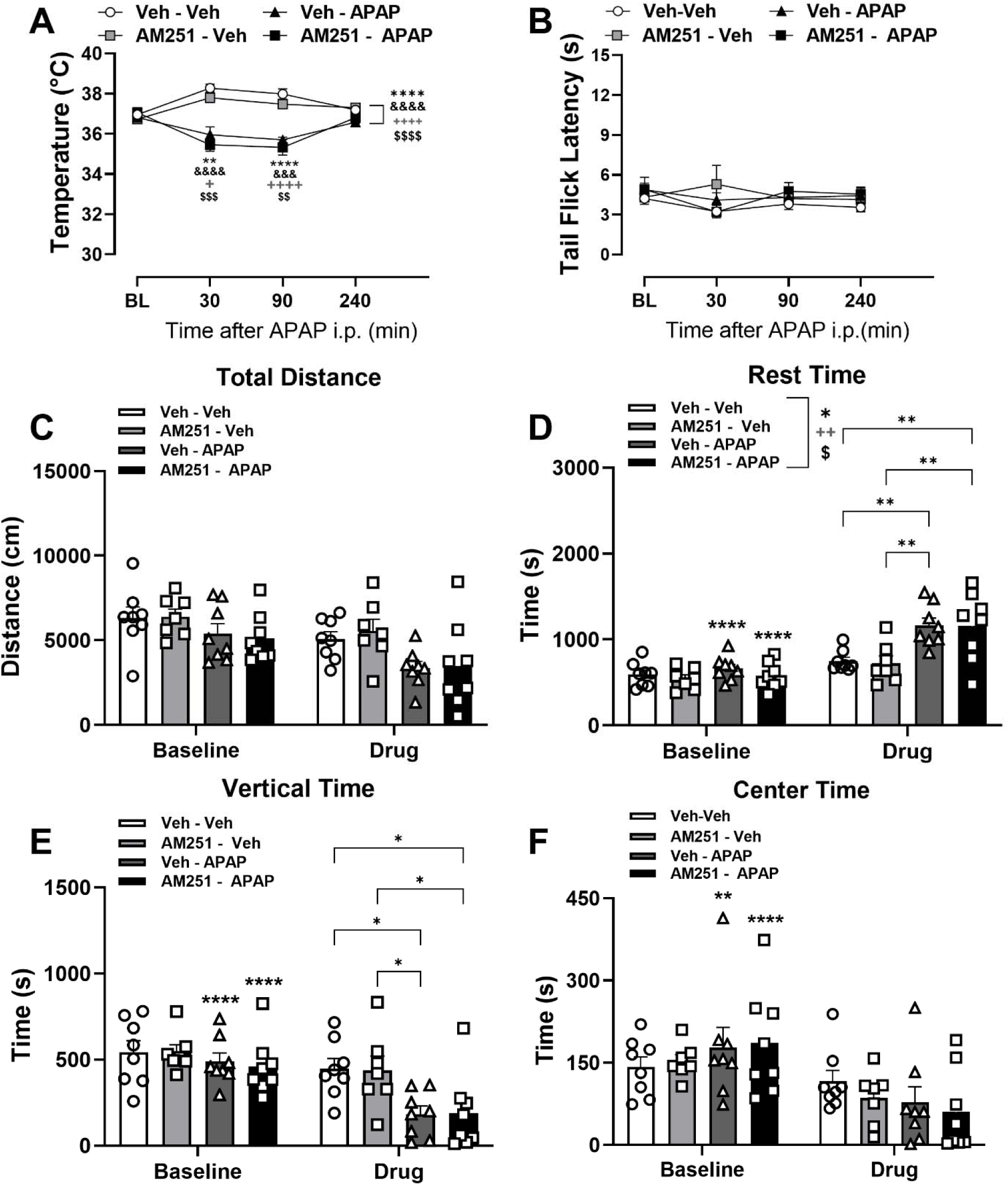
The CB1 receptor antagonist (5 mg/kg, i.p.) does not block effects of APAP (300 mg/kg, i.p.) on body temperature or locomotion. (**A**) Body temperature of naïve male mice after treatment with APAP. (**B**) Tail flick latency after treatment with APAP. (**C-F**) Activity meter data reflecting locomotion (center time, rest time, vertical time, and total distance) 30 minutes after treatment with APAP (n=7-8 per group). Data show mean (± SEM). Two-way ANOVA followed by Bonferroni’s post hoc test. Interaction between time points and treatment effect (Panels A and B): *\*\* P<0.01, **** P <0.0001 Veh-Veh vs. Veh-APAP; + P<0.05, ++++ P<0.0001 AM251-Veh vs Veh-APAP; &&& P<0.001, &&&& P<0.0001 Veh-Veh vs. AM251-APAP; $$ P<0.01, $$$ P<0.001 AM251-Veh vs. AM251-APAP. Brackets (main treatment effect only, Panel A-F): * P <0.05, **** P <0.0001 Veh-Veh vs. Veh-APAP; ++ P<0.01, ++++ P<0.0001 AM251-Veh vs. Veh-APAP; & P<0.05, &&&& P<0.0001 Veh-Veh vs. AM251-APAP; $ P<0.05, $$$$ P<0.0001 AM251-Veh vs. AM251-APAP. Lines between bars (interaction treatment and time, Panel C-F): * P <0.05, ** P <0.01, *** P <0.001 group difference. Symbols on top of group bars at baseline (Panel C-F): * P <0.05, ** P <0.01, *** P <0.001, **** P <0.0001 difference from same group at drug administration time point*.

APAP increased rest time independent of CB1 receptors (**Figure 3D**, Treatment: F_3,27_=4.366, p=0.0125; Interaction: F_3,27_=7.727, p=0.0007). Rest time was elevated overall in Veh-APAP compared to Veh-Veh (p=0.0142) and Veh-AM251 (p=0.0045) treatment, and in AM251-APAP relative to AM251-Veh (p=0.0249) treatment. Veh-APAP increased post-injection levels of rest time relative to Veh-Veh (p=0.0019) and AM251-Veh (p=0.0013) and did not differ from AM251-APAP (p> 0.9999) treatment. AM251-APAP elevated rest time compared to all control conditions (vs. Veh-Veh, p=0.0023, vs. AM251-Veh, p=0.0016). Accordingly, rest time was reduced compared to baseline numbers in all mice given APAP (Veh-APAP p<0.0001; AM251-APAP p<0.0001).

APAP also reduced vertical time in the activity meter independent of CB1 (**Figure 3E**, Treatment: F_3,27_=2.550, p=0.0767; Interaction: F_3,27_=6.037, p=0.0028). Veh-APAP reduced vertical time relative to Veh-Veh (p=0.0229) and AM251-Veh (p=0.0409) and did not differ from AM251-APAP (p> 0.9999) treatment. AM251-APAP elevated rest time compared to all control conditions (vs. Veh-Veh, p=0.0283, vs. AM251-Veh, p=0.0499). In comparison to pre-injection baselines, APAP also reduced vertical time irrespective of pre-treatment with AM251 (Veh-APAP p<0.0001; AM251-APAP p<0.0001).

APAP, administered alone or following AM251 pre-treatment, did not alter time spent in the center of the activity meter overall but the interaction between treatment and time was significant (**Figure 3F**, Treatment: F_3,27_=0.0356, p=0.9908; Interaction: F_3,27_=3.045, p=0.0459). Irrespective of AM251 pretreatment, APAP decreased center time compared to respective pre-injection (baseline) levels (Veh-APAP p=0.0016; AM251-APAP p<0.0001), but center time was similar between groups assessed after pharmacological manipulations. Baseline responding in the activity meter were not different between groups in this subset of experiments for any of the parameters mentioned above.

Lastly, APAP (100 mg/kg i.p.) did not alter body temperature (**Supplemental Figure 3A**), tail-flick latency (**Supplemental Figure 3B**), distance traveled, rest time, vertical time or center time in the activity meter apparatus (**Supplemental Figure 3C-F**).

### 3.6 DAGL inhibitor RHC-80267 does not attenuate cannabimimetic effects of APAP

In male mice, APAP (300 mg/kg, i.p.) reduced body temperature overall and in a time-dependent manner but the DAGL inhibitor RHC-80267 (20 mg/kg) did not block these effects (**Figure 4A**, Treatment: F_3,25_=9.418, p=0.0002; Interaction: F_9,75_=12.92, p<0.0001); both Veh-APAP and RHC-APAP produced equivalent (p>0.9999) reductions in body temperature relative to either Veh-Veh or RHC-Veh treatment (p<0.0001 for each comparison). Effects of RHC-80267-Veh did not differ from Veh-Veh ((p>0.9999). APAP did not alter tail flick latencies in otherwise naïve male mice (**Figure 4B**).

**Figure 4.**
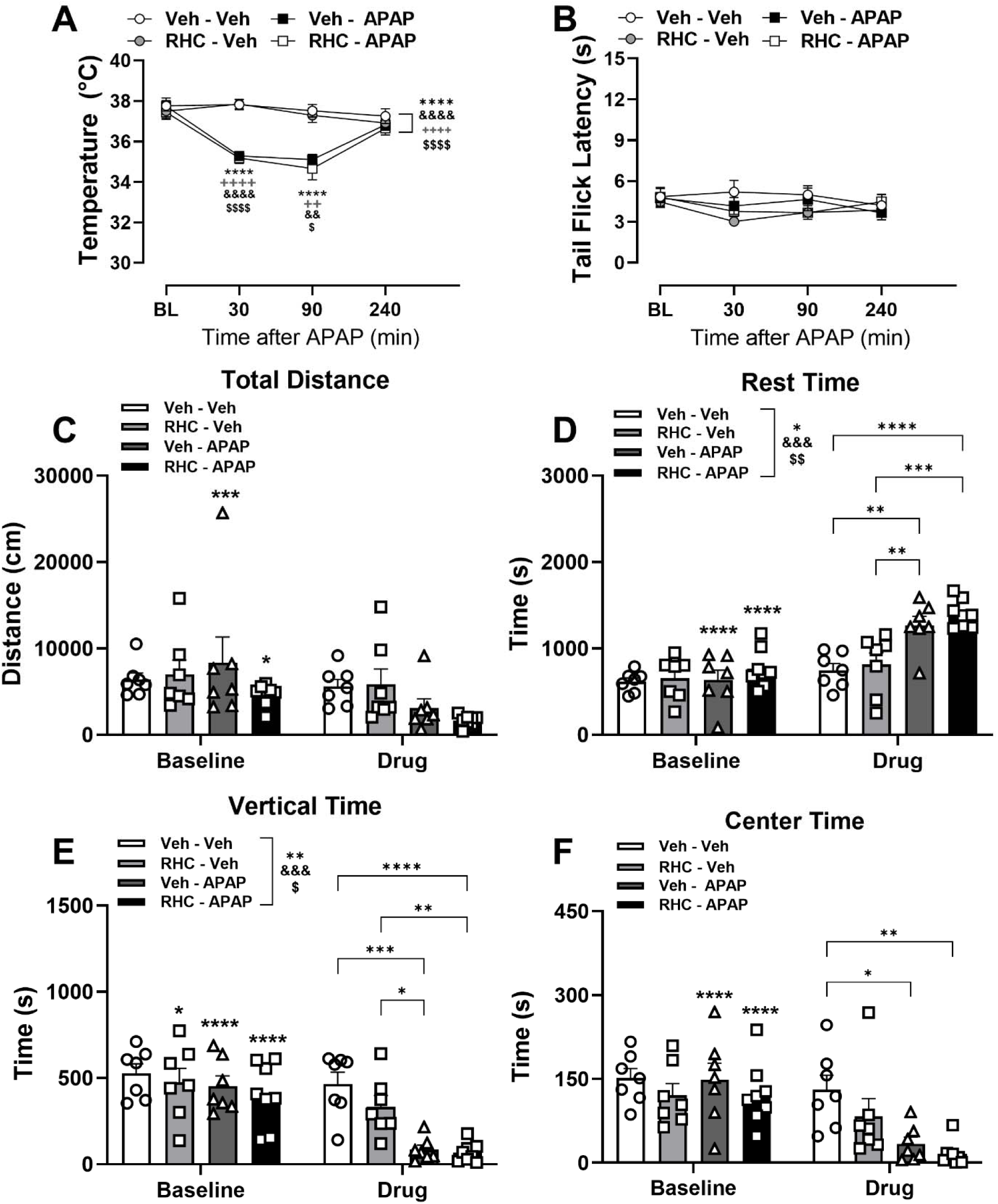
APAP (300 mg/kg, i.p.) produces hypothermia and hypolocomotion, but not tail flick antinociception in otherwise naïve male mice. The DAGL inhibitor RHC-80267 (20 mg/kg) does not block the side effects induced by APAP. (**A**) Body temperature of naïve male mice after treatment with Veh-Veh, Veh-APAP, RHC80267-Veh or RHC80267-APAP. (**B**) (**C-F**) Activity meter data reflecting locomotion (center time, rest time, vertical time, and total distance) 30 minutes after pharmacological manipulations (n=7-8 per group). Data show mean (± SEM). Two-way ANOVA followed by Bonferroni’s post hoc test. Interaction between time points and treatment effect (Panels A and B): *\*\*\*\* P <0.0001 Veh-Veh vs. Veh-APAP; ++ P<0.01, ++++ P<0.0001 RHC-Veh vs Veh-APAP; && P<0.01, &&&& P<0.0001 Veh-Veh vs. RHC-APAP; $ P<0.05, $$$$ P<0.0001 RHC-Veh vs. RHC-APAP. Brackets (main treatment effect only, Panel A-F): * P <0.05, ** P <0.01, **** P <0.0001 Veh-Veh vs. Veh-APAP; ++++ P<0.0001 RHC-Veh vs. Veh-APAP; & P<0.05, && P<0.01, &&& P<0.001, &&&& P<0.0001 Veh-Veh vs. RHC-APAP; $ P<0.05, $$ P<0.01, $$$$ P<0.0001 RHC-Veh vs. RHC-APAP. Lines between bars (interaction treatment and time, Panel C-F): * P <0.05, ** P <0.01, *** P <0.001, **** P <0.0001 group difference. Symbols on top of group bars at baseline (Panel C-F): * P <0.05, ** P <0.01, *** P <0.001, **** P <0.0001 difference from same group at drug administration time point*.

Activity meter parameters did not differ between groups before drug treatment (P>0.4297). Drug treatments altered total distance traveled in a time-dependent manner (**Figure 4C**; Treatment: F_3,25_=1.277, p=0.3038; Interaction: F_3,25_=3.858, p=0.0214), but post hoc comparisons failed to reveal between-group differences post-injection (p>0.2457). APAP reduced total distance from baseline regardless of RHC-80267 pretreatment (Veh-APAP: p=0.0001; RHC-APAP: p=0.0183). Overall, activity meter parameters varied over time (p<0.05), and showed significant main effects of treatment and interaction between factors (see **Supplemental Table 1**).

APAP increased rest time but these effects were not blocked by RHC80267 (**Figure 4D**; Treatment: F_3, 25_ = 5.032, p=0.0073; Interaction: F_3,25_ =19.87, p<0.0001). Veh-APAP did not differ from RHC-APAP (p>0.9999) treatment but increased post-injection rest time relative to Veh-Veh (p=0.0013) and RHC-Veh (p=0.0062) controls. Rest time in RHC-Veh groups did not differ from Veh-Veh treatment (p>0.9999). APAP increased rest time from baseline in both groups (p<0.0001).

APAP also decreased vertical time overall and these effects were not blocked by RHC80267 (**Figure 4E**; Treatment: F_3, 25_ =5.571, p=0.0046; Interaction: F_3, 25_=8.639, p=0.0004). Veh-APAP reduced vertical time compared to Veh-Veh (p=0.0002) and RHC-Veh (p=0.0098) treatments that did not differ from each other (p>0.9999). RHC-APAP also reduced vertical time relative to Veh-Veh (p<0.0001) or RHC-Veh (p= 0.0098) but did not differ from Veh-APAP treatment (p>0.9999). All groups except Veh-Veh showed reduced vertical time from baseline (p<0.0410).

Similarly, APAP reduced center time post-injection (**Figure 4F**; Treatment: F_3, 25_=2.559, p=0.0777; Interaction: F_3, 25_=5.062, p=0.0071). Both Veh-APAP (p=0.0190) and RHC-APAP (p=0.0023) reduced center time relative to Veh-Veh and their own baselines (p<0.0001).

### 3.7 APAP attenuated mechanical hypersensitivity but not paw edema in CFA-injected mice

In male mice, APAP (30-300 mg/kg, i.p.) produced a dose- and time-dependent increase in paw withdrawal thresholds in the CFA-injected (ipsilateral) paw (**Figure 5A**: Treatment: F_3,22_=16.43, p<0.0001; Interaction: F_9,66_=3.474, p=0.0015). APAP at 100 (p=0.0037) and 300 mg/kg (i.p.) (p<0.0001) increased ipsilateral paw withdrawal in comparison to the vehicle-treated group (**Figure 5A**). The high dose increased withdrawal thresholds overall to a greater extent than the low dose (p=0.0208).

**Figure 5.**
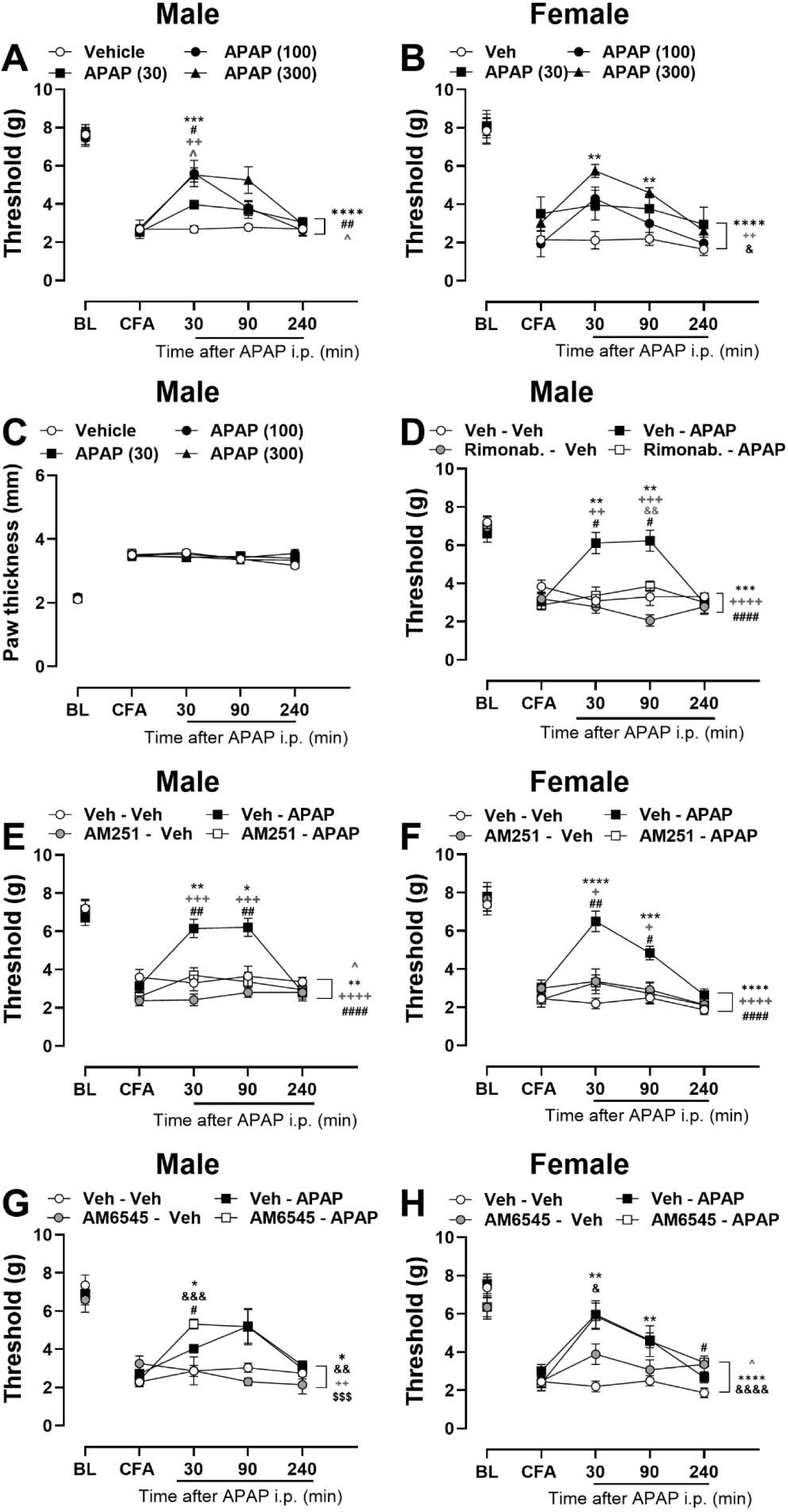
APAP dose-dependently suppresses CFA-induced mechanical hypersensitivity without altering paw edema in mice and the anti-allodynic efficacy is blocked by global, but not peripherally restricted CB1 receptor antagonists. APAP elevated mechanical paw withdrawal threshold in the CFA-injected (ipsilateral) paw of (**A**) male and (**B**) female mice at different doses. APAP did not alter thickness of the paw (**C**) ipsilateral to CFA injection in male mice. CB1 receptor antagonist Rimonabant (5 mg/kg, i.p.) suppresses the anti-allodynic effect of APAP (300 mg/kg, i.p.) evaluated in the ipsilateral paw of male mice with CFA-induced inflammatory pain (**D**). CB1 receptor antagonist AM251 (5 mg/kg, i.p.) (**E, F**), but not AM6545 (10 mg/kg, i.p.) (**G, H**) suppress the anti-allodynic effect of APAP (300 mg/kg, i.p.) evaluated in the ipsilateral paw of male (**E, G**) and female (**F, H**) mice with CFA-induced inflammatory pain (n=5-9 per group). ***Panel A-C****: Data show mean (± SEM) for ipsilateral paws. Two-way ANOVA followed by Bonferroni’s post hoc test. Interaction between time points and treatment effect: ** P <0.01, *** P <0.001 Vehicle vs. APAP 300; # P <0.05 Vehicle vs. APAP 100; ++ P<0.01 Vehicle vs. APAP 30; ^ P<0.05 APAP 30 VS. APAP 300. Brackets (main treatment effect only): **** P <0.0001 Vehicle vs. APAP 300; ## P <0.01 Vehicle vs. APAP 100; ++ P<0.01 Vehicle vs. APAP 30; ^ P<0.05 APAP 30 VS. APAP 300. & P<0.05 APAP 100 vs. APAP 300. **Panel D**: Data show mean (± SEM). Two-way ANOVA followed by Bonferroni’s post hoc test. Interaction between time points and treatment effect: ** P <0.01, Veh-Veh vs. Veh-APAP; # P <0.05, Veh-APAP vs. Rimonab-APAP; ++ P <0.01, +++ P<0.001 Rimonab-Veh vs. Veh-APAP; && p<0.01 Rimonab-Veh vs Rimonab-APAP. Brackets (main treatment effect only): *** P <0.001 Veh-Veh vs. Veh-APAP; #### P <0.0001 Veh-APAP vs. Rimonab-APAP; ++++ P<0.0001 Rimonab-Veh vs. Veh-APAP. **Panel E-H**:* Data show mean (± SEM). Two-way ANOVA followed by Bonferroni’s post hoc test. Interaction between time points and treatment effect: *\* P<0.05, ** P <0.01, *** P <0.001, **** P <0.0001 Veh-Veh vs. Veh-APAP; #P <0.05, ## P <0.01, Veh-APAP vs. AM251-APAP or Veh-APAP vs. AM6545-APAP; +P <0.05, +++ P<0.001 AM251-Veh vs. Veh-APAP or AM6545-Veh vs. Veh-APAP; & P<0.05, &&& P<0.001 Veh-Veh vs. AM6545-APAP. Brackets (main treatment effect only): ** P <0.01, **** P <0.0001 Veh-Veh vs. Veh-APAP; ## P <0.01, ### P <0.001, #### P <0.0001 Veh-APAP vs. AM251-APAP; ++++P<0.0001 AM251-Veh vs. Veh-APAP; ^ P<0.05 Veh-Veh vs. AM251-Veh; && P<0.01, &&&& P<0.0001 Veh-Veh vs. AM6545-APAP; $$$ P<0.001 AM6545-Veh vs. AM6545-APAP. ^ P<0.05 Veh-Veh vs. AM6545-Veh*.

In female mice, APAP at 30 (p=0.0028) and 300 mg/kg (p<0.0001) increased mechanical thresholds in the CFA-injected paw relative to vehicle treatment in a dose and time-dependent manner (**Figure 5B**; Treatment: F_3,16_=3.384, p=0.0442; Interaction: F_9,48_=2.522, p=0.0187); the 300 mg/kg dose produced greater anti-allodynic efficacy that the 100 mg/kg dose. Pharmacological treatments did not alter contralateral paw withdrawal thresholds (**Supplemental Figure 4B**).

APAP did not alter paw thickness at any dose in either the ipsilateral (**Figure 5C**; Treatment: F_3,22_=0.3116, p=0.8168; Interaction: F_9,66_=1.338, p=0.2347) or contralateral (**Supplemental Figure 4C**; Treatment: F_3,20_=2.441, p=0.0942; Interaction: F_9,60_=1.673, p=0.1157) paw of male mice.

### 3.8 Global, but not peripherally restricted CB1 receptor antagonists attenuate the effect of APAP on CFA-induced inflammatory pain in mice

A structurally distinct CB1 antagonist, Rimonabant, blocked the anti-allodynic effect of APAP (300 mg/kg i.p.) in male mice (**Figure 5D**; Treatment: F_3,28_=14.42, p<0.0001; Interaction: F_9,84_=7.180, p<0.0001); Veh-APAP (300 mg/kg) elevated paw withdrawal thresholds in comparison to Veh-Veh, Rimonabant-Veh or Rimonabant-APAP (p<0.0001 for each comparison) treatments, which did not differ from each other (p≥0.0690). None of the treatments altered withdrawal thresholds in the contralateral paw (**Supplemental Figure 4D**; Treatment: F_3,28_=0.1319, p=0.9403; Interaction: F_9,84_=1.349, p=0.2247).

The CB1 antagonist AM251 blocked the anti-allodynic effect of APAP in male mice (**Figure 5E**; Treatment: F_3,27_=12.40, p<0.0001; Interaction: F_9,81_=6.107, p<0.0001). Veh-AM251, by itself, did not alter thresholds of the CFA-injected (ipsilateral) paw compared to pre-injection levels (F_3,27_=1.774, p=0.1759). Withdrawal thresholds were lower in the AM251-Veh compared to Veh-Veh group (p=0.0222) but did not differ from the post-CFA threshold determined prior to pharmacological manipulations (p≥0.61 for all timepoints). Veh-APAP increased ipsilateral paw withdrawal thresholds relative to Veh-Veh (p=0.0021) or AM251-Veh (p<0.0001) groups and AM251-APAP (p<0.0001) treatment.

In female mice, APAP increased thresholds in CFA-injected paw in a CB1-dependent manner (**Figure 5F**; Treatment: F_3,27_=14.98, p<0.0001; Interaction: F_9,81_=4.850, p<0.0001). Veh-APAP increased ipsilateral paw thresholds relative to Veh-Veh, AM251-Veh (p<0.0001) groups and AM251-APAP (p<0.0001) treatment.

In male mice, the peripherally-restricted CB1 antagonist AM6545 (10 mg/kg) did not attenuate the anti-allodynic effect of APAP (**Figure 5G**; Treatment: F_3,20_=5.833, p=0.0049; Interaction: F_9,60_=3.641, p=0.0011). Veh-APAP increased withdrawal thresholds in the CFA-injected paw overall compared to Veh-Veh (p=0.0150) or AM6545-Veh (p=0.0064) treatments that did not differ from each other (p>0.9999). AM6545 did not attenuate the anti-allodynic effect of APAP (p>0.9999).

In female mice, APAP increased paw withdrawal thresholds in the CFA-injected paw irrespective of AM6545 pre-treatment (**Figure 5H**; Treatment: F_3,26_=12.39, p<0.0001; Interaction: F_9,78_=3.538, p=0.0010); Veh-APAP and AM6545-APAP increased withdrawal thresholds relative to Veh-Veh (p< 0.0001) treatment and effects of AM6545-APAP did not differ from Veh-APAP overall (p>0.9999). Paw withdrawal thresholds were reliably elevated overall in AM6545-Veh compared to the Veh-Veh group (p=0.0380) but did not differ from post-CFA pre-injection thresholds at any post-injection timepoint (p≥0.1546).

Neither AM251 nor AM6545, in the presence or absence of APAP, altered contralateral paw withdrawal thresholds in either sex (**Supplemental Figure 4E-H**).

### 3.9 DAGL and MAGL inhibitors attenuate the anti-allodynic effect of APAP in CFA-induced inflammatory pain

In male mice, APAP (300 mg/kg (i.p.) given after vehicle pre-treatment (Veh-APAP), increased paw withdrawal thresholds in the CFA-injected paw (**Figure 6A**; Treatment: F_3,24_=12.27, p<0.0001; Interaction: F_9,72_=4.183, p=0.0002) in comparison to either Veh-Veh (p<0.0001) or RHC-Veh (p<0.0001) and RHC-80267 pretreatment blocked the anti-allodynic effect of APAP (p=0.0021).

**Figure 6.**
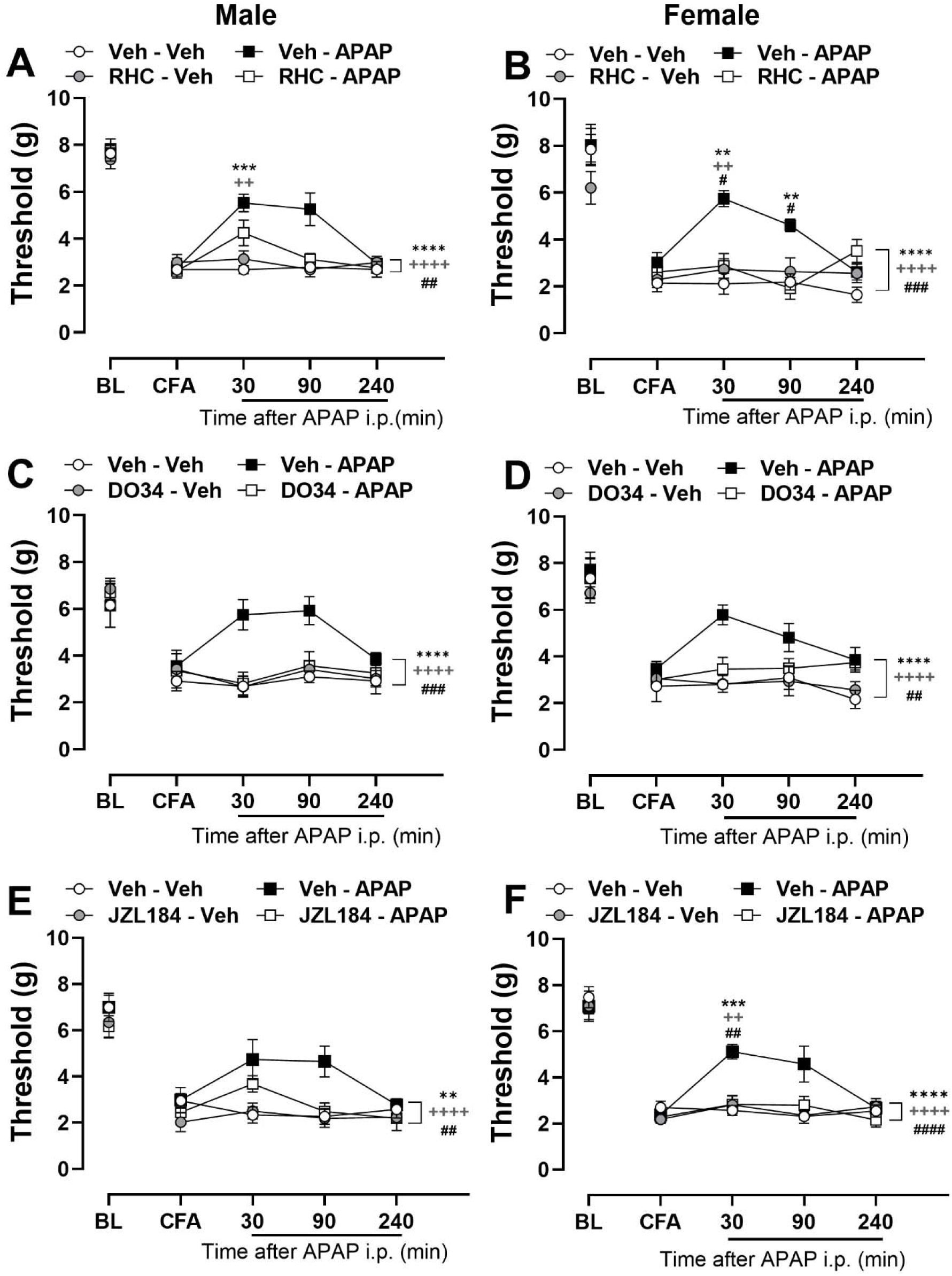
Anti-allodynic effects of APAP are prevented by pharmacological inhibitors of DAGL and MAGL in a mouse model of inflammatory pain. DAGL inhibitor RHC-80267 (20 mg/kg, i.p.) attenuates the anti-allodynic effect of APAP (300 mg/kg, i.p.) in the CFA-injected (ipsilateral) paw of (**A**) male and (**B**) female mice. DAGL inhibitor DO34 (30 mg/kg, i.p.) attenuates the anti-allodynic effect of APAP (300 mg/kg, i.p.) in the CFA-injected (ipsilateral) paw of (**C**) male and (**D**) female mice. MAGL inhibitor JZL 184 (4 mg/kg, i.p.) attenuates the anti-allodynic effect of APAP (300 mg/kg, i.p.) in the CFA-injected (ipsilateral) paw of (**E**) male and (**F**) female mice. Data show mean (± SEM) (n = 5-7 per group). Two-way ANOVA followed by Bonferroni’s *post hoc* test. Interaction between time points and treatment effect: *\*\* P <0.01, *** P <0.001 Veh-Veh vs. Veh-APAP; # P <0.05 Veh-APAP vs. RHC-APAP or Veh-APAP vs. DO34-APAP; ## P <0.01 Veh-APAP vs. JZL 184-APAP; ++ P<0.01 RHC-Veh vs. Veh-APAP or DO34-Veh vs. Veh-APAP or JZL184-Veh vs. Veh-APAP. Brackets (main treatment effect only):**** P <0.0001 Veh-Veh vs. Veh-APAP; ## P <0.01, ### P <0.001, #### P <0.0001 Veh-APAP vs. RHC-APAP or Veh-APAP vs. DO34-APAP or Veh-APAP vs. JZL 184-APAP; ++++ P<0.0001 RHC-Veh vs. Veh-APAP or DO34-Veh vs. Veh-APAP or JZL 184-Veh vs. Veh-APAP* .

In female mice, APAP increased paw withdrawal thresholds in the CFA-injected paw in a DAGL -dependent manner (**Figure 6B**; Treatment: F_3,16_=19.49, p<0.0001; Interaction: F_9,48_=3.813, p=0.0011), Veh-APAP (300 mg/kg, i.p.) increased ipsilateral paw withdrawal thresholds relative to Veh-Veh or RHC-Veh treatment (p<0.0001 for each comparison) and RHC80267 blocked the anti-allodynic effect of APAP (p=0.0003).

In male mice, Veh-APAP (300 mg/kg, i.p.) increased paw withdrawal thresholds in the CFA-injected paw in a DAGL-dependent manner (**Figure 6C**; Treatment: F_3,20_=9.825, p=0.0003; Interaction: F_9,60_=1.887, p=0.0713). Veh-APAP increased ipsilateral paw withdrawal thresholdds compared to either Veh-Veh (p<0.0001) or DO34 (30 mg/kg)-Veh alone (p<0.0001) and DO34 blocked the anti-allodynic effect of APAP (p=0.0003).

In female mice, APAP elevated paw withdrawal thresholds in the CFA-injected paw in a DAGL-dependent manner (**Figure 6D**; Treatment: F_3,19_=11.39, p=0.0002; Interaction: F_9,57_=1.564, p=0.1485). Veh-APAP (300 mg/kg, i.p.) increased thresholds in the CFA-injected paw relative to Veh-Veh (p<0.0001) or DO34-Veh (p<0.0001) groups. DO34 pretreatment blocked the anti-allodynic effect of APAP (p=0.0071).

In male mice, APAP attenuated CFA-induced mechanical hypersensitivity in a MAGL-dependent manner. Veh-APAP (300 mg/kg) increased thresholds in the CFA-injected mice compared to JZL-APAP (p=0.0071), Veh-Veh (p=0.0022) or JZL184-Veh (p<0.0001) groups (**Figure 6E**; Treatment: F_3,17_=8.080, p=0.0015; Interaction: F_7.667,43.45_=1.681, p=0.1333).

In female mice, APAP increased paw withdrawal thresholds in the CFA-injected paw in a MAGL-dependent manner (**Figure 6F**; Treatment: F_3,20_=7.967, p=0.0011; Interaction: F_6.430,42.87_=4.229, p=0.0016). Veh-APAP increased thresholds compared to Veh-Veh and JZL184-Veh groups and this anti-allodynic effect of APAP was blocked by JZL184 (p<0.0001 for each comparison). Thus, pre-treatment with JZL184 blocked the anti-allodynic effect of APAP (300 mg/kg, i.p.) in mice of both sexes.

In general, pharmacological treatments did not reliably alter paw withdrawal thresholds in the non-inflamed paw in either male or female mice (**Supplementary 5A-F**). A single main effect of treatment was observed in contralateral paw withdrawal thresholds in male mice receiving JZL184 (**Supplementary Figure 5E**; Treatment: F_3,17_=4.085, p=0.0235; Interaction: F_7.022,39.79_=1.166, p=0.3439), but *post hoc* analysis did not reveal differences at any time point compared to the baseline injection (p>0.05).

### 3.10 APAP treatment modulates endogenous lipids levels differentially across the CNS

To evaluate the broader range of lipid signaling modulation beyond the established reduction in prostaglandins induced by APAP, an exploratory study was conducted in which 99 endogenous small-molecule lipids were screened. The total numbers detected at analytical levels in each tissue (i.e., having a minimum of 4 samples with a chromatographic peak above the signal to noise ratio for that endogenous lipid) was as follows: spinal cord (SC), 72; rostral ventral medulla (RVM), 56; periaqueductal gray (PAG), 64, thalamus (THAL), 63; and paw skin, 74. Of those endogenous lipids detected at analytical levels in each CNS region, the following percentage of lipids changed in each structure as follows: SC, 24%, RVM, 32%, PAG, 39%, THAL, 14%, and PS, 20%. **Figures 7** and **8** illustrate specific findings on the direction and magnitudes of these changes.

**Figure 7.**
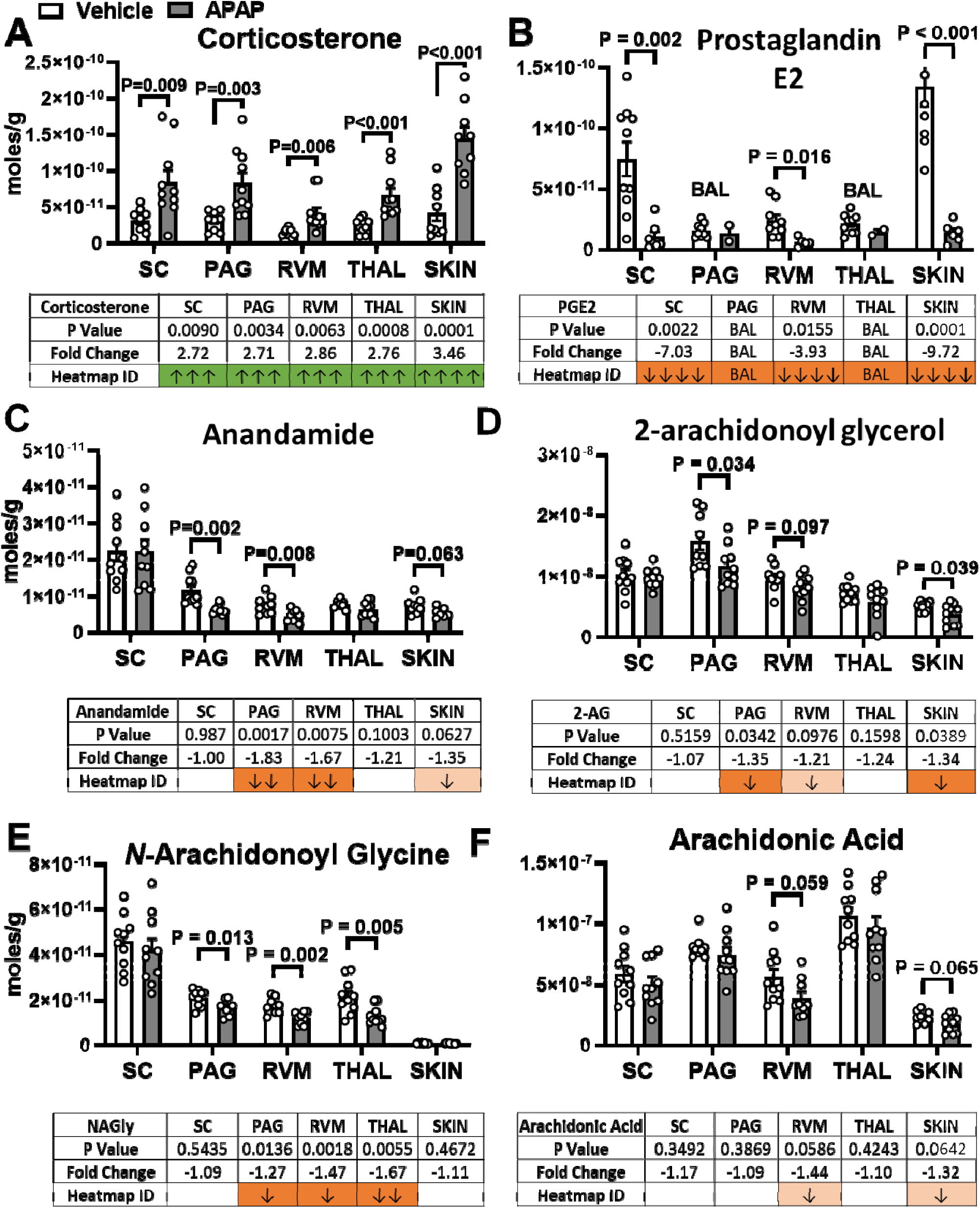
Representative bar graphs of endogenous lipids in 4 CNS tissue and paw skin (SKIN) with heatmap icons. The bar graphs on the on the top of the examples are the mean SEM of the levels of endogenous lipids in treatment group (Control in White and IMQ treatment in Blue). (**A**) Corticosterone (Cort); (**B**) Prostaglandin E2 (PGE2); (**C**) Arachidonoyl ethanolamine (AEA; Anandamide); (**D**) 2-arachidonoyl glycerol (2-AG); (**E**) Arachidonoyl glycine (NAGly); (**F**) Arachidonic acid (AA). The table underneath the bar graphs list the analytical data of the APAP verses vehicle comparisons. See Methods for description of Python data analysis for Fold change (denoted by arrows) and statistics (colors denote significance levels and direction) to generate the Heatmap colors and arrows. Data illustrate the means, SEM, and individual data points in bar graphs for each of 6 endogenous lipids in the vehicle and APAP treatment groups.

**Figure 8.**
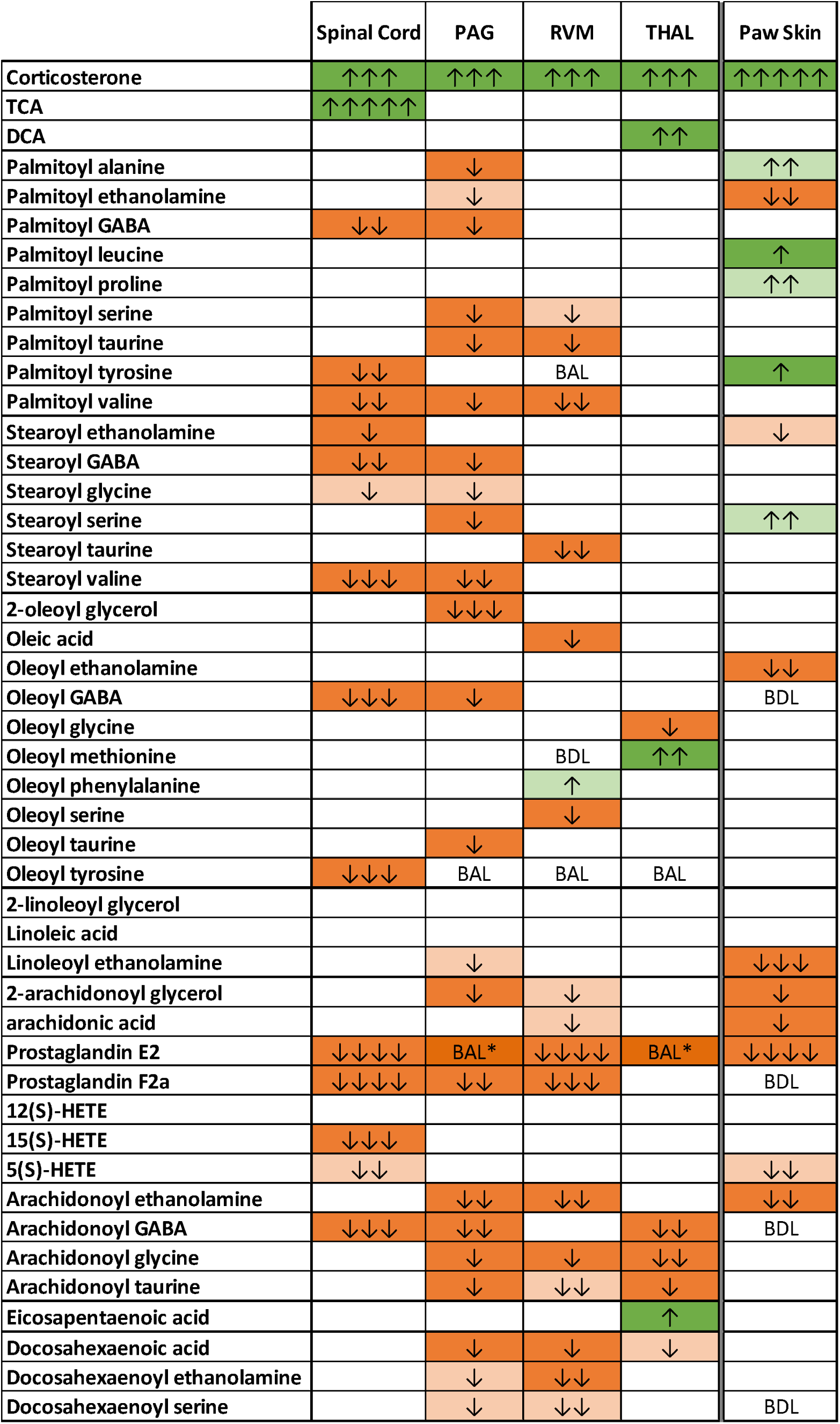
Heatmap of endogenous lipids detected at analytical levels in CNS and paw skin tissue. Dark green represents significant increases after treatment (p ≤0.05) while light green represents significance levels of p ≤ 0.1-0 >.05. Dark orange represents significant decreases after treatment (p ≤0.05) while light orange represents significance levels of p ≤ 0.1-0 >.05. Fold change is indicated by the number of arrows, where 1 arrow corresponds to 1–1.49-fold difference, 2 arrows to a 1.5–1.99-fold difference, 3 arrows to a 2–2.99-fold difference, 4 arrows a 3–9.99-fold difference, and 5 arrows a difference of tenfold or more. BAL (below analytical limits) refers to endogenous lipids that were measured in at least one sample but less than 4 making statistical analysis unreliable. BDL (below detectable limits) refers to endogenous lipids that were not detected in any samples. Blank cells indicate that endogenous lipids were present in at least 4 samples in each treatment group being analyzed and that no significant differences were measured.

Endogenous lipids were significantly modulated by APAP treatment (see **Figure 7** for example). Graphical representation of the data, p-values, fold-change, and heatmap identifier are shown in **Figure 8** (see Methods for more details on heatmaps). Analysis for all 99 endogenous lipids screened are shown in **Supplemental Figure 6**. Corticosterone levels increased 2.7-2.9-fold across all CNS tissues and 3.1-fold in the paw skin (Figure 7A). The only other endogenous lipid that showed significant changes across all tissues was PGE2 (**Figure 7B**). APAP decreased PGE2 in SC, RVM, and paw skin with the levels being reduced below detection thresholds in the PAG and THAL. Levels of AEA (**Figure 7C**) and 2-AG (**Figure 7D**) were reliably decreased in PAG, RVM, and skin and levels of AA (**Figure 7F**) were only reduced in RVM and paw skin. Importantly, the AEA metabolite, *N*-arachidonoyl glycine (NAGly; **Figure 7E**) were reliably reduced in PAG, RVM, and THAL.

A selected heatmap of endogenous lipids that showed significant changes with APAP treatment is shown in **Figure 8**. The full heatmap for all endogenous lipids is provided in **Supplemental Figure 6**. The heatmap is organized by acyl group with those cholesterol-based lipids (e.g., corticosterone and bile acids) at the top, followed by classes in order of acyl chain length and level of desaturation (e.g., docosahexaenoic, C22:6; arachidonic, C20:4; linoleic, C18:2; oleic, C18:0; stearic, C18:1; and palmitic, C16:0). Three patterns emerged in the data. 1) The overall effect of APAP in the CNS is a decrease in endogenous lipid levels, with the important exceptions of corticosterone and bile acids and palmitoyl conjugates in paw skin. 2) PAG and SC exhibited the most similarity in overall changes, 3) arachidonic acid derivatives had the highest percentage of specific lipids species modulated within the acyl chain category in the CNS and palmitic acid derivatives had the highest percentage change in paw skin.

### 3.11 APAP is highly abundant but the metabolite AM404 is not detectible in CNS tissue

Chromatograms and quantitative data for the levels of APAP and its putative metabolite, AM404, in the CNS are shown in **Figure 9**. The levels of APAP in all tissues were in the range of 1-2 micro moles/gram which was calculated from the final calculated molarity of the lipid extract that was 0.9nM in this example (**Figure 9A**). The levels of APAP in the non-diluted sample were close to the ceiling of the analytical detection level, which made the values less reliable, therefore, the sample was diluted 1:100 to more accurately evaluate APAP levels. APAP levels in the extract were ∼2-fold higher than the endogenous levels of 2-AG measured there and 10-1000-fold higher than most endogenous lipids in the current exploratory lipidomics screen. The chromatogram of a 10 µl injection of the 1:100 dilution of a spinal cord sample with a calculated concentration of 0.9 nM being equivalent to the peak height and area to 10 fmols of APAP, which is the result of a 10 µl injection of a 1 nM (1E-9) APAP standard is shown in **Figure 9A**. The signal to noise ratio for APAP detection here is ∼1 fmol. The detection limits of AM404, ∼2.5 fmols, with 5 and 10 µl injections of 1nM AM404 (i.e., 5 and 10 fmol respectively) are shown in **Figure 9B**. However, no AM404 was detected in any brain region with injections up to 40ul of non-diluted samples (**Figure 9B**). The levels of APAP and AM404 across all 4 CNS regions are summarized in **Figure 9**.

**Figure 9.**
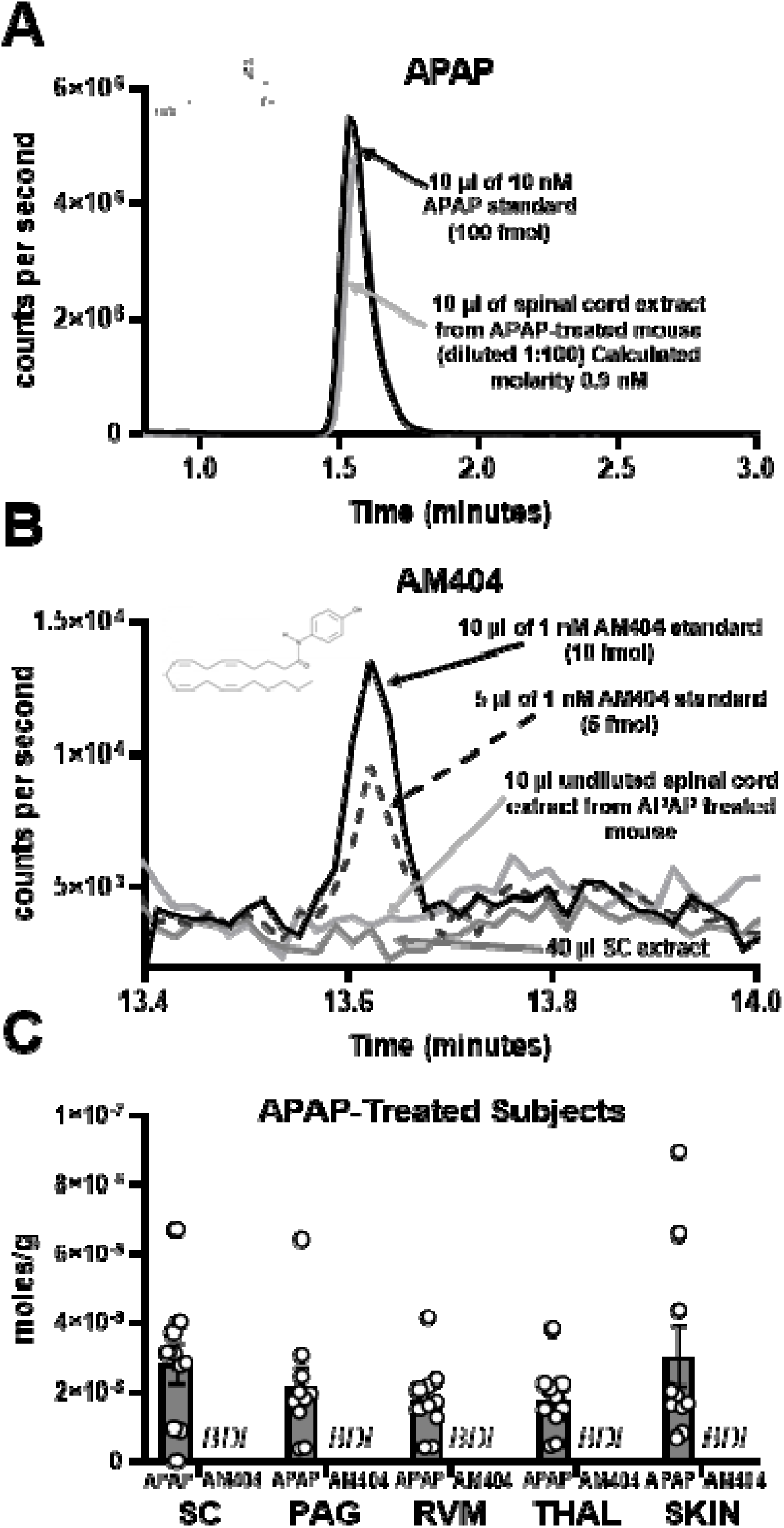
APAP, but not AM404, is detected in CNS and paw skin (SKIN) tissue via HPLC/MS/MS. (**A**) Overlaid chromatograms of APAP standard (black line; 152.11/110.02) and spinal cord (SC) extract diluted 1:100 (gray line; 152.11/110.02). APAP molecular structure cartoon inset for reference. (**B**) Overlaid chromatograms of AM404 standard (black lines; 396.33/152.02 and SC extract 1:1 (gray line; 396.33/152.02). AM404 molecular structure cartoon inset for reference. (**C**) Mean (± SEM) for APAP and AM404 in SC, periaqueductal gray (PAG), rostral ventromedial medulla (RVM), thalamus (THAL) and paw skin (SKIN). Individual data points in open circles. Samples derived from CFA-injected mice treated with APAP (300 mg/kg, i.p.).

The pharmacological mechanisms underlying APAP-induced analgesia across mechanistically distinct preclinical pain models and in mice of both sexes are shown in **Figure 10**.

**Figure 10.**
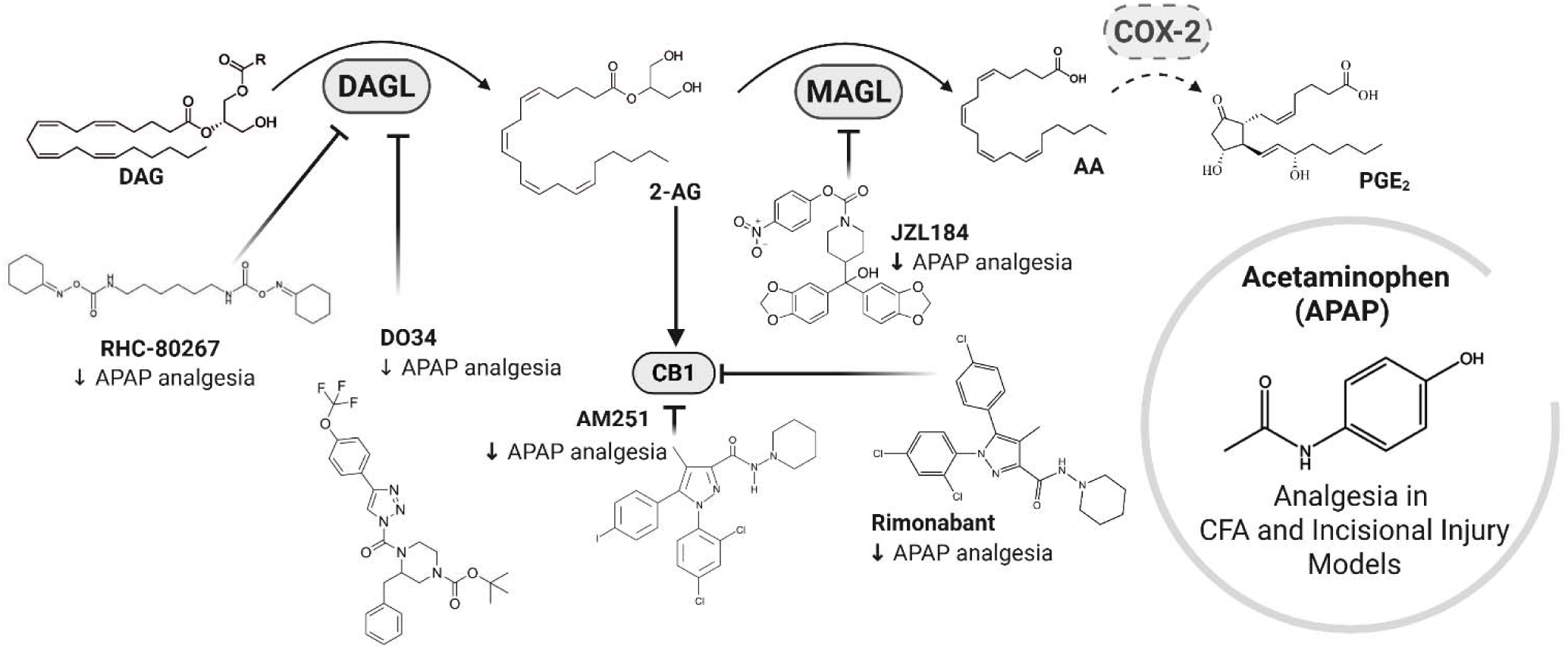
Graphical summary of the pharmacological findings from pain behavior assays in APAP-induced analgesia in CFA and Incisional Injury Models in mice. 2-arachidonoylglycerol (2-AG); Acetaminophen (APAP); Arachidonic acid (AA); Complete Freund’s Adjuvant (CFA); Cyclooxygenase-2 (COX-2); Diacylglycerol (DAG); Diacylglycerol lipase (DAGL); Monoacylglycerol lipase (MAGL); Prostaglandin E2 (PGE2).

## 4. Discussion

Here we show that APAP attenuates post-surgical and inflammatory pain behavior in mice through a mechanism that requires the enzymes DAGL and MAGL, as well as CB1 cannabinoid receptors. These conclusions are validated using mechanistically distinct pathological pain models, both sexes, two mouse strains and different routes of administration (p.o. and i.p.). Furthermore, these effects are unlikely to be attributed to the metabolism of APAP to AM404, as AM404 was not detected in either CNS or paw skin samples at the peak of APAP-induced analgesia. No changes in mechanical thresholds in the uninjured paw were observed in any study. Thus, APAP-induced elevations in paw withdrawal thresholds required a pathological pain state. APAP suppressed CFA-induced mechanical hypersensitivity without altering paw edema, consistent with prior work suggesting that systemic administration of APAP lacks a peripheral anti-inflammatory property [31–33]. Nonetheless, APAP treatment significantly reduced PGE2 within portions of the CNS of CFA-injected mice (Spinal cord, RVM), and in paw skin tissue. In addition, APAP increased corticosterone levels across tissue from CNS, as well as paw skin, and reduced endogenous lipids such as 2-AG and AEA (PAG, RVM and paw skin).

Initial investigations of APAP-induced antinociception have mainly focused on tests of acute nociception (e.g., hot plate test, paw pressure test) [17,34–36], that have uncertain translational relevance. Investigations of APAP’s analgesic mechanisms require testing in translationally relevant pre-clinical models of pathological pain. Here, APAP suppressed mechanical hypersensitivity induced by both incisional injury and intraplantar CFA at doses that did not alter spinally-mediated tail-flick antinociception (see [37]). APAP (300 mg/kg i.p.) increased rest time, and decreased vertical activity, without altering total distance traveled, and reduced body temperature. APAP is reported to exert thermoregulatory properties through inhibition of cyclooxygenase, leading to reduced production of pyrogenic prostaglandins in hypothalamus and reduction of fever (for review see [38]). However, the effects on locomotor activity and body temperature were not mediated by CB1 cannabinoid receptors or DAGL and cannot impact interpretation of mechanisms of APAP-induced anti-allodynic efficacy observed here.

Microinjection of APAP metabolites, but not APAP itself, into the ventrolateral PAG decreases pain thresholds, suggesting that APAP must be metabolized to act within the CNS [18]. Previous reports suggest that APAP’s analgesic effects are partly mediated by its central conversion to AM404, a bioactive metabolite that modulates pain signaling via TRPV1 and CB1 receptor activation [39–41]. However, in our study, AM404 was not detected in any of the CNS tissue (spinal cord, PAG, RVM, thalamus) or paw skin tissue taken 90 minutes post APAP treatment in CFA-injected mice. Previous pharmacokinetic and human cerebrospinal fluid (CSF) studies have reported that AM404 is present at very low concentrations under clinically relevant APAP dosing concentrations. For instance, brain levels in rodents are shown in picogram-per-gram range with minimal detectability, while human CSF concentrations are found below established pharmacological activity thresholds [42,43]. These findings have led to ongoing debate regarding whether sufficient AM404 is generated *in vivo* to meaningfully contribute to APAP’s analgesic effects [44]. However, a limitation of our study is that our lipidomic analysis performed in tissue dissected at 90 minutes post APAP treatment does not fully exclude the possibility of transient, localized, or earlier formation of AM404 [42].

Previous studies have demonstrated the involvement of CB1 receptors in the antinociceptive effects of APAP in *in vivo* models of inflammatory pain [16,17]. In addition, deletion of CB1 cannabinoid receptors has previously demonstrated to increase prostaglandins (PGE2, PGF2α) in several portions of the mouse brain [27]. Here, the antinociceptive efficacy of APAP was abolished by CNS-penetrant CB1 receptor antagonist/inverse agonists (AM251, Rimonabant), but not by the peripherally restricted antagonist AM6545. AM251 per se did not affect body temperature or locomotor activity in naïve mice, suggesting that the off-target effects [45–48] observed *in vitro* are unlikely to explain our data.

Our findings are the first to report that structurally distinct DAGL inhibitors (RHC-80267 and DO34), at doses that did not themselves alter nociceptive thresholds, attenuated the anti-allodynic effect of APAP in two different models of pathological pain and in mice of both sexes. In addition, a behaviorally inactive dose of JZL184, a MAGL inhibitor, attenuated the anti-allodynic effect of APAP in both the CFA and incisional injury models. Our pharmacological data suggests that 2-AG metabolism and blocking of CB1 receptors may interfere with the analgesic efficacy of APAP. However, 2-AG levels were not elevated in our samples 90 minutes post APAP injection. More work is necessary to determine whether 2-AG, which is known to inhibit cyclooxygenase-2 (COX-2) [49], could be elevated at earlier time points.

2-AG is one of the main sources of arachidonic acid (AA) in the brain [50,51]. Regulation of 2-AG release is associated with the suppression of pain signaling through DAGLα and CB1-dependent mechanisms [26]. However, 2-AG synthesis or degradation may not result in measurable changes in 2-AG levels within pain-related neural circuits of *ex vivo* tissue [52]. 2-AG signaling in microenvironments may also be masked by larger pools of 2-AG in dissected tissues (see also [26,53]) given its role as an intermediate in lipid metabolism. Our behavioral findings suggest that APAP’s mechanism is tied to regulation of endogenous lipids in the DAGL and MAGL axis, presumably 2-AG (see also [26]); however, the lipidomic analysis did not support this hypothesis. 2-AG was decreased or unaltered, and little to no change in AA was observed at 90 minutes post injection. A transient spike in 2-AG could potentially trigger inhibition of COX-2 and decrease prostaglandin production from AA [49], although anti-allodynic efficacy of APAP was still prominent at this time point. Inhibition of COX-2 can enhance retrograde endocannabinoid signaling, which indicates COX-2 normally acts to limit endocannabinoid-mediated synaptic suppression [54]. Notably, transient decreases in 2-AG levels results from chronic stress hormone exposure [55], which may help explain our lipidome results at 90 minutes from tissue of CFA-injected mice given APAP. Alternately, 2-AG may not be a key signaling molecule in APAP signaling.

APAP treatment in CFA-injected mice produced >2 fold levels of corticosterone (analog of human cortisol) across brain, spinal cord and paw skin tissue and decreased prostaglandins in spinal cord, RVM and paw skin. APAP exposure can influence stress-related signaling pathways and oxidative balance in the body [56,57]. In fact, increased levels of steroid hormones (e.g., cortisol) driven by administration of APAP have been reported in humans [58–62]. High cortisol levels have been directly associated with high paracetamol levels in hair samples of young adults [58]. APAP also inhibits CYP17A1 enzyme activity and limits glucocorticoid release in human adrenocortical cells [59], which could lead to adrenocorticoid hormone release *in vivo*, favoring the upregulation of other steroid hormones (e.g., corticosterone and aldosterone) [63,64]. Peripheral inflammation induced by CFA can increase systemic levels of corticosterone, but results are inconsistent [59,65,66], and corticosterone levels may rapidly subside [67]. In addition, APAP-induced liver injury in mice is positively correlated with increased corticosterone levels [67], and prevented by glucocorticoid receptor inhibitor administration in mice [57]. Elevated corticosterone levels during stress have been shown to suppress central prostaglandin synthesis via glucocorticoid-mediated inhibition of cyclooxygenase pathways [57,68,69], which may contribute to the analgesic efficacy of APAP (300 mg/kg, i.p.) by dampening prostaglandin-dependent signaling [70]. Corticosterone can regulate inflammation and prostaglandin production (e.g., PGE2) [70,71], and administration of exogenous corticosteroids reliably attenuates inflammatory pain and edema (e.g., carrageenan, and CFA-induced inflammatory pain models and neuropathic pain models) [72–76]. Further studies are required to investigate the role of glucocorticoid receptor activation in potentially mediating the effects of APAP on analgesia and/or toxicity.

Stress exposure, stress hormones and exogenous glucocorticoids can exert complex effects on lipid signaling, including endocannabinoid signaling (see [65,77]). Glucocorticoids can rapidly mobilize endocannabinoids, which act on CB1 receptors to suppress excitatory neurotransmission [78,79]. Modulation of endogenous lipids by glucocorticoids depends on both the intensity and duration of stress (see [65]). Although APAP treatment is not inherently a stressor, it can modulate the hypothalamic-pituitary-adrenal axis [58,59] and, consequently, alter endocannabinoid levels in tissue. AEA and other endogenous lipids such as palmitoyl ethanolamine (PEA) are known activators of the ion channel TRPV1 [80,81], which can lead to increase excitatory neurotransmission and sensitization in the periphery (e.g., intraplantar injection in rodents; primary sensory neurons; pro-nociceptive effect) [82,83]. However, after APAP treatment, corticosterone levels were increased across CNS and paw skin tissue, whereas AEA and 2-AG levels were decreased in PAG, RVM and paw skin of CFA-injected mice. Further studies are needed to evaluate the time course of endogenous lipid modulation in tissue after APAP treatment, particularly at early time points [53] and the potential role of TRPV1 receptors in the antinociceptive effects of APAP [39] in pathological pain models.

Our results suggest that pharmacological inhibition of DAGL and MAGL, as well as glucocorticoid modulation by APAP, may contribute to wide-scale endogenous lipid signaling modulated by APAP administration. Our lipidomic findings may not fully capture early transient signaling events, compartmentalized lipid pools, or changes in lipid turnover [13,52,84]. However, up to 39% of the targeted lipids in the screen were modulated in the CNS at 90 minutes, which presents a completely heretofore unrecognized endogenous lipid modulation caused by APAP treatment. Given the dramatic decrease in PGE2 with little effect on AA and 2-AG at the 90-minute time point, there is a particular interest in three additional AA-derived endogenous lipids; *N*-arachidonoyl glycine (NAGly), N-arachidonoyl GABA (NAGABA); and *N*-arachidonoyl taurine (NA-taurine). Each of these endogenous signaling lipids was reduced with a greater effect size than AEA or 2-AG and in more CNS regions. NAGABA is the least studied of the 3, yet it’s first introduction in 2001 by Michael Walker’s group showed it was involved in pain processing [85] but recent data shows that it is involved in cerebrovascular activity that is associated with PGE2 [86]. NA-taurine is the second most studied with a wider range of signaling properties with the earliest identification of NA-taurine showing activity at TRP channels by Ben Cravatt’s group [87,88]. More recently, NA-taurine was shown to be elevated in hepatic tissue in metabolic dysfunction associated steatotic liver disease (MASLD; [89]). In that the level of APAP used in this study that causes analgesia is also used to cause liver toxicity, there may be a relationship in which endogenous signaling/metabolic pathways are involved with APAP and its role in corticosterone, liver function, and regulation of endogenous lipids. Finally, NAGly has been more broadly studied and has signaling properties immune function and inflammation in part through its modulation of GPR18 [90–93].

DAGL and MAGL are ubiquitous enzymes in the body and brain [94,95] and are likely differentially modulated by drugs such as APAP, or their respective pharmacological inhibitors (e.g., RHC-80267, DO34, JZL184), which also means that their effects on endogenous lipid signaling would be tissue-dependent and is what we report here. Further research is required to determine how CB1 receptors are involved in these endogenous lipid signaling pathways, given our findings showing and overall reduction endogenous lipid signaling following APAP treatment and not an increase in endocannabinoids. Earlier work from our group showed that constitutive CB1 knockout caused significant elevations in resting PGE2 across all CNS tissues [96]. Lipidomic analysis of Rimonabant showed significant decreases in triglycerides, and cholesterol esters but only a moderate reduction in phosphatidyl ethanolamines; however, no endocannabinoids or other endogenous lipid signaling molecules were measured [97]. Taken together; however, there is evidence that both constitutive and acute inhibition of CB1 can significantly change endogenous lipids.

In conclusion, APAP-induced analgesia is associated with central CB1 receptors, as well as enzymes implicated in the biosynthesis and metabolism of a broad range of signaling lipids. APAP treatment produced marked increase in corticosterone levels, a key modulator of inflammatory mediators and endogenous signaling lipid pathways. Importantly, our studies demonstrate that the analgesic and side-effect profiles of APAP are mechanistically distinct.

Moreover, our targeted pharmacological and exploratory lipidomic findings support a previously unrecognized mechanism underlying APAP-induced analgesia.

## Acknowledgments

We thank Macie DeLillo, Jasmin Bola, and Elyse Chaffee for helping with mass spectrometric analysis.

---

**Supplemental Figure 1.**
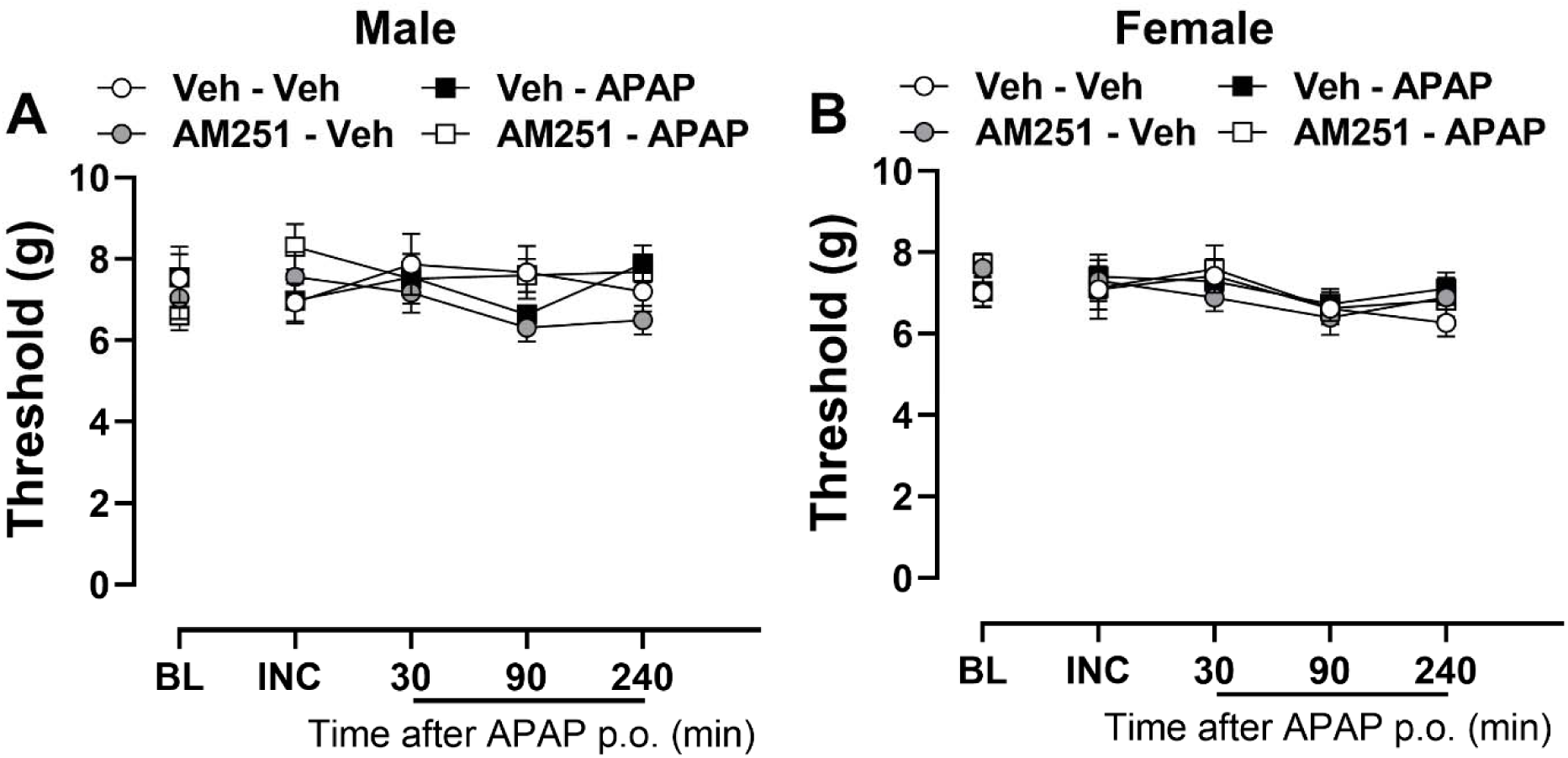
AM251 (5 mg/kg, i.p.) and APAP (300 mg/kg, p.o.) do not alter mechanical paw withdrawal thresholds evaluated in the contralateral paw of male (**A**) and female mice (**B**) with incisional injury (n= 6 per group). Data show mean (± SEM) for contralateral paws. Two-way ANOVA followed by Bonferroni’s post hoc test.

**Supplemental Figure 2.**
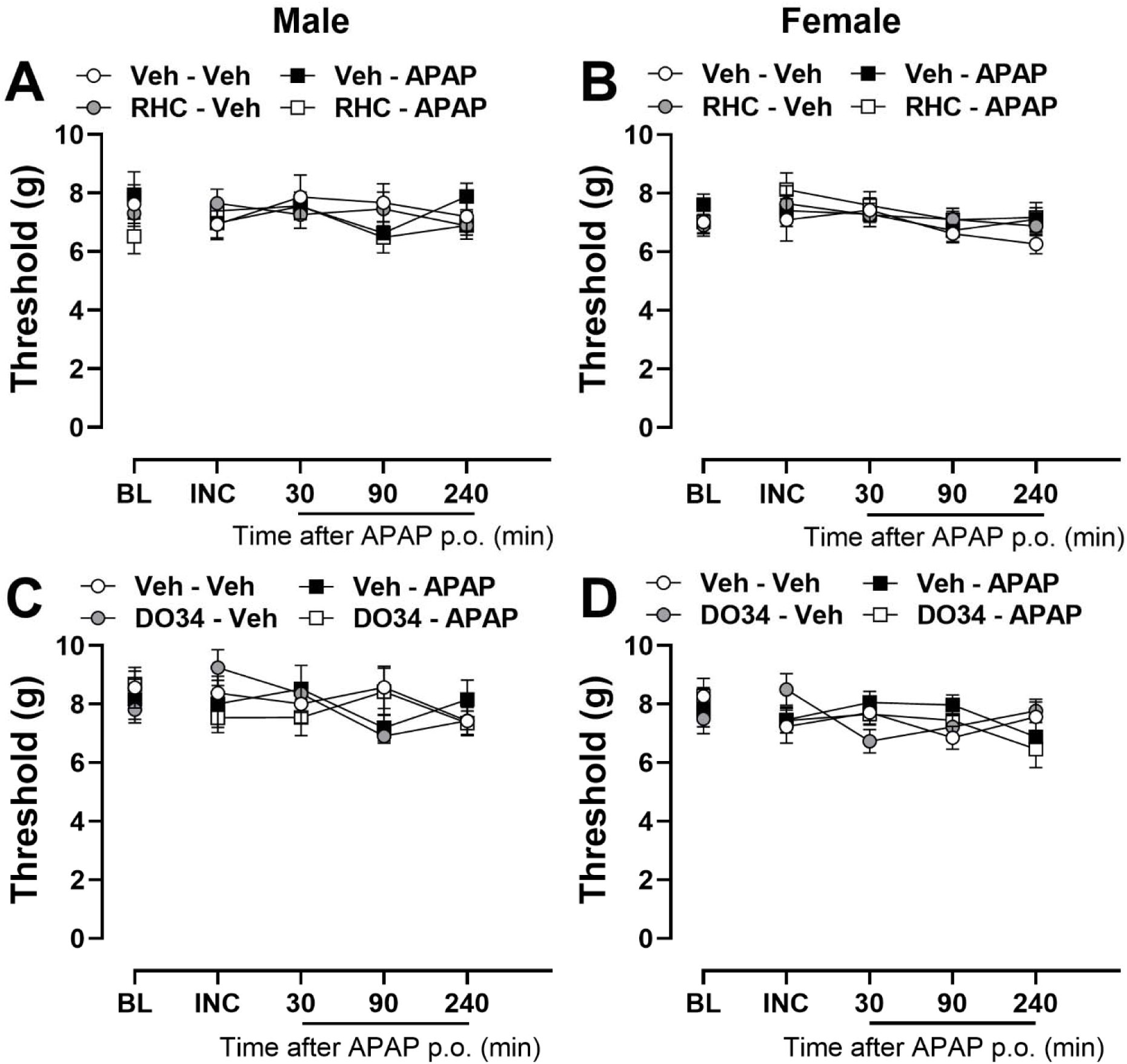
DAGL inhibitors RHC-80267 (20 mg/kg, i.p.) (**A, B**) or DO34 (30 mg/kg, i.p.) (**C, D**), and APAP (300 mg/kg, p.o.) do not alter mechanical paw withdrawal thresholds evaluated in the contralateral paw of male and female mice with incisional injury (n=6 per group). Data show mean (± SEM) for contralateral paws. Two-way ANOVA followed by Bonferroni’s post hoc test.

**Supplemental Figure 3.**
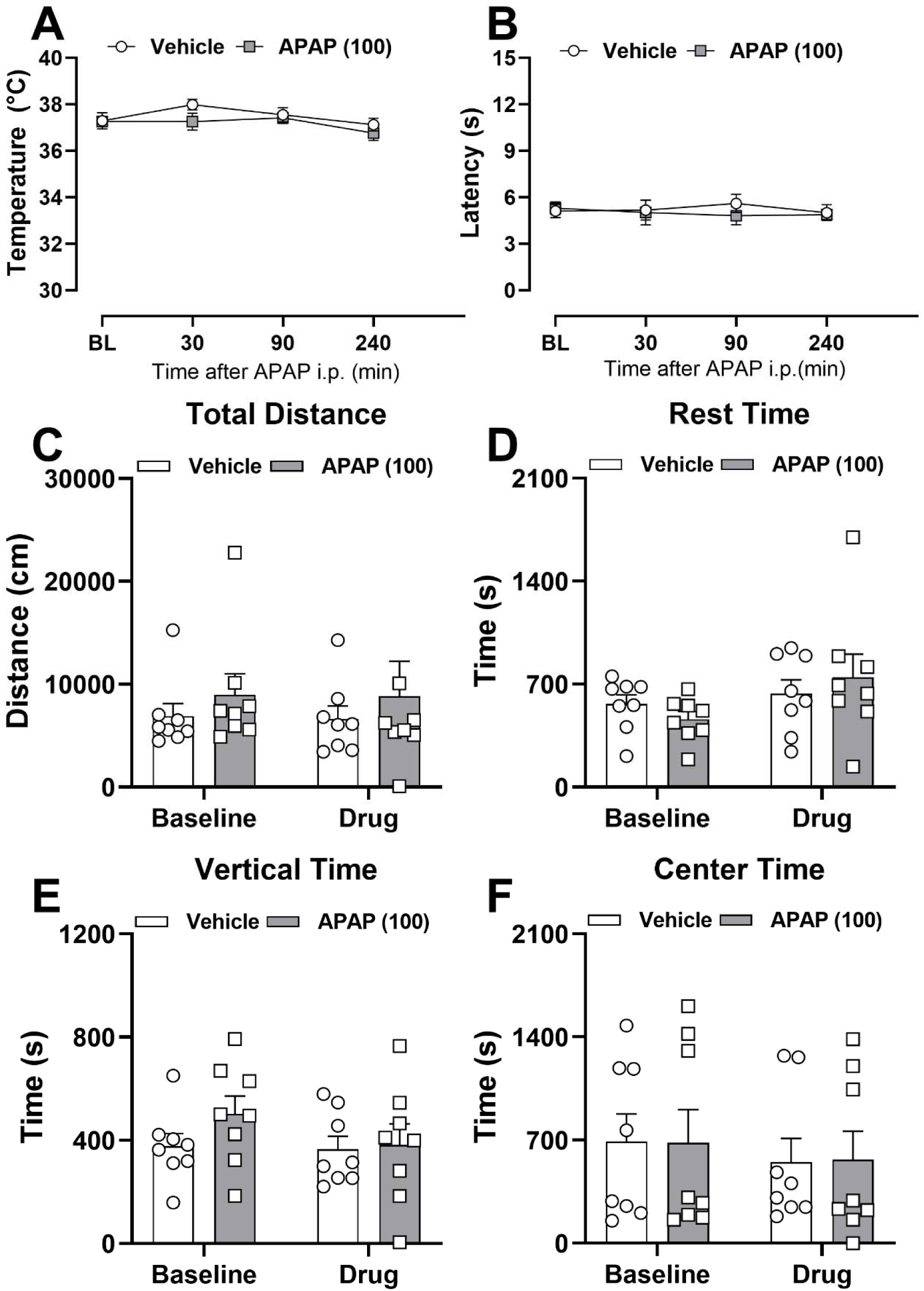
APAP (100 mg/kg, i.p.) does not produce hypothermia, hypolocomotion or tail flick antinociception in naïve male mice. (**A**) Body temperature of naïve male mice after treatment with APAP 100 mg/kg. (**B**) Tail flick latency after treatment with APAP 100 mg/kg. (**C-F**) Activity meter data reflecting locomotion (center time, rest time, vertical time, and total distance) 30 minutes after treatment with APAP 100 mg/kg (n=8 per group). Data show mean (± SEM). Two-way ANOVA followed by Bonferroni’s post hoc test.

**Supplemental Figure 4.**
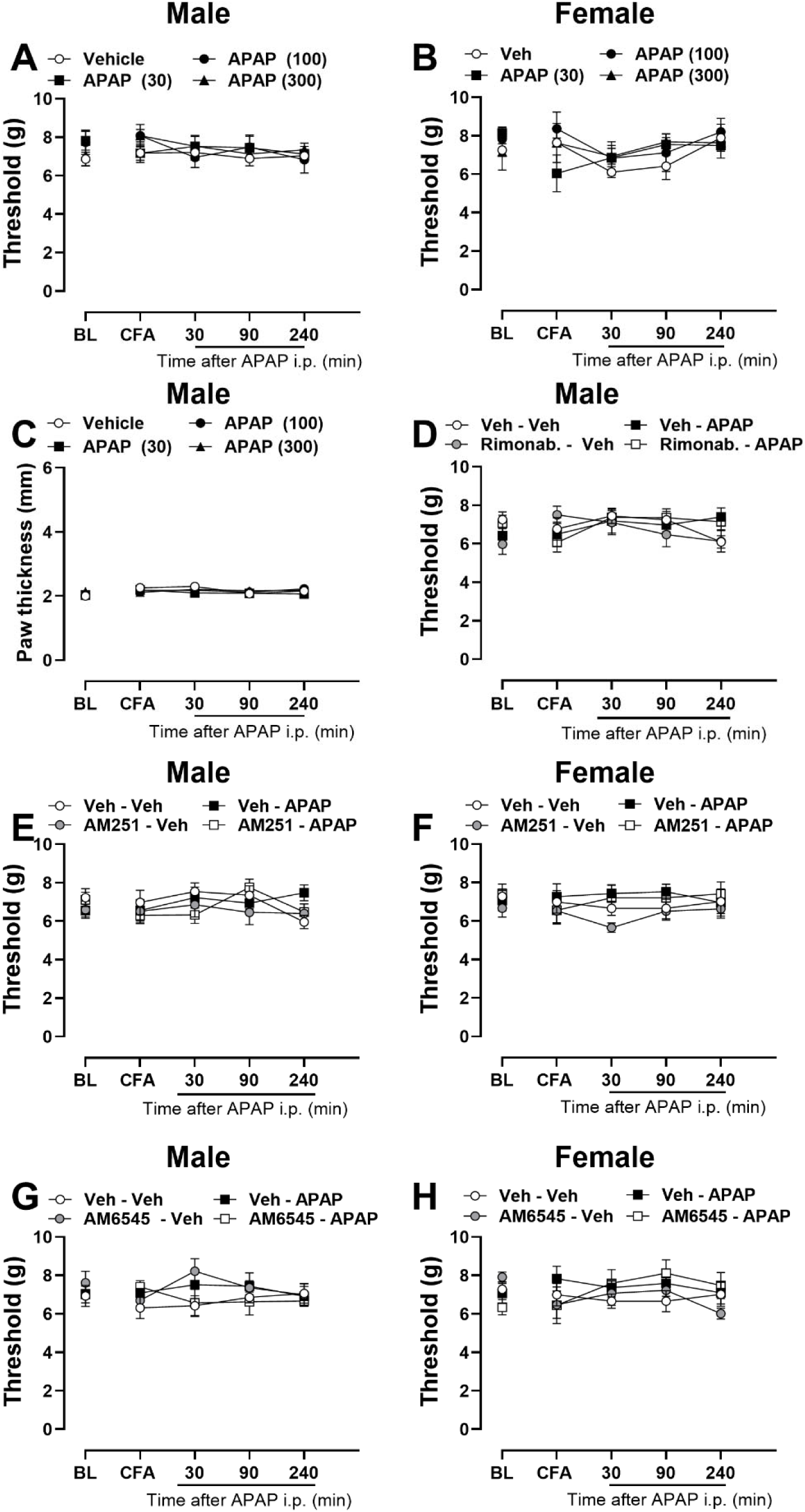
APAP does not affect the withdrawal threshold of the contralateral paw following CFA injection in (**A**) male and (**B**) female mice and does not alter paw thickness in (**C**) CFA-injected male mice. The CB1 receptor antagonist Rimonabant (5 mg/kg, i.p.) does not change the withdrawal threshold of the contralateral paw in male mice (**D**). Similarly, the CB1 receptor antagonist AM251 ( 5 mg/kg, i.p.) does not affect the contralateral paw withdrawal threshold in (**E**) male or (**F**) female mice. The peripherally restricted CB1 receptor antagonist AM6545 (10 mg/kg, i.p.) also does not alter mechanical thresholds of the contralateral paw in (**G**) male or (**H**) female mice. Data are presented as mean ± SEM (n = 5–9 per group). Statistical analysis was performed using two-way ANOVA followed by Bonferroni’s post hoc test.

**Supplemental Figure 5.**
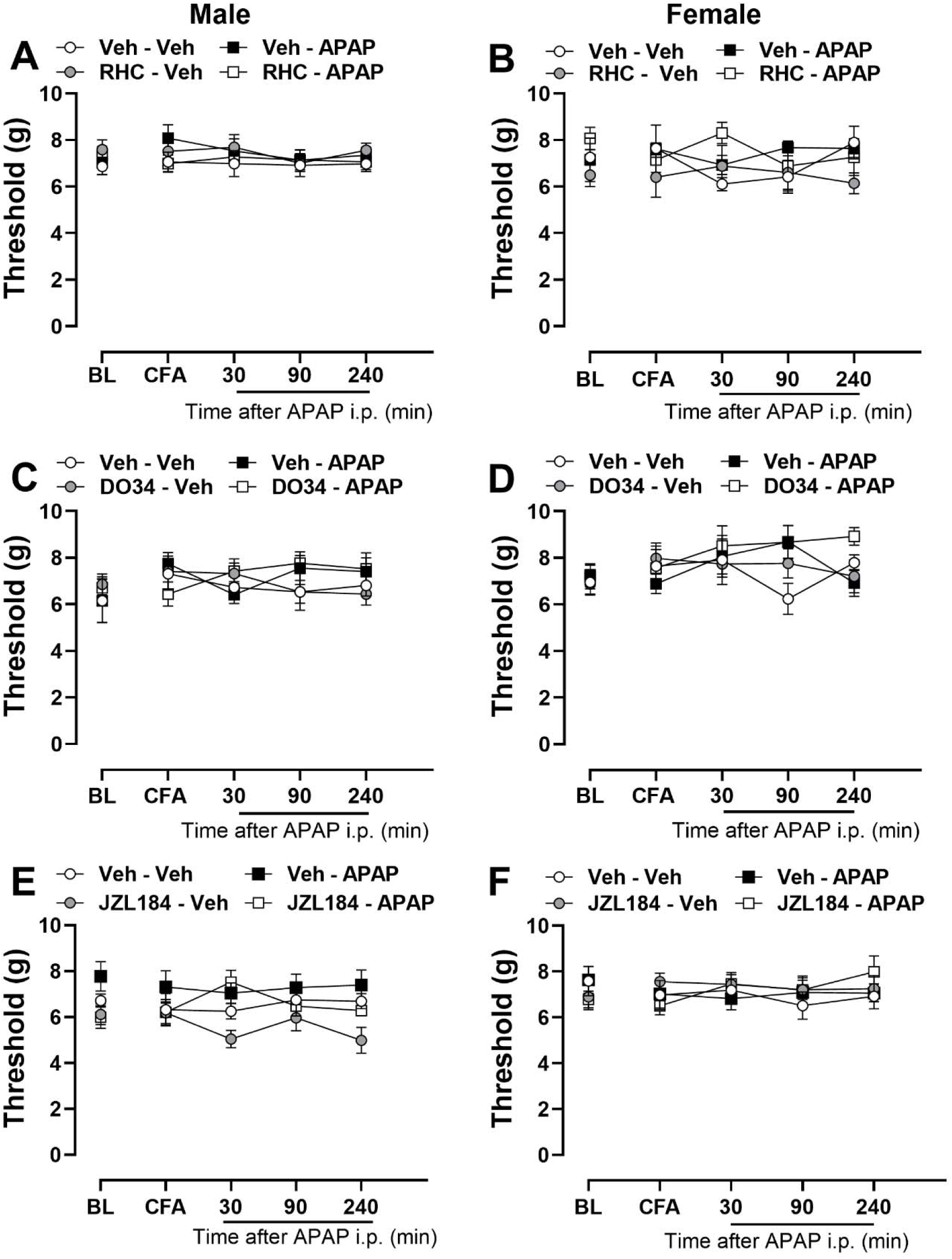
DAGL inhibitor RHC-80267 (20 mg/kg, i.p.) (**A, B**) or DO34 (30 mg/kg, i.p.) (**C, D**), or MAGL inhibitor JZL184 (1.4 mg/kg, i.p) (**E, F**) and APAP (300 mg/kg, i.p.) do not alter mechanical paw withdrawal thresholds evaluated in the contralateral paw of male and female mice with CFA-induced inflammatory pain (n= 5-7 per group). Data show mean (± SEM) for contralateral paws. Two-way ANOVA followed by Bonferroni’s post hoc test.

**Supplemental Figure 6.**
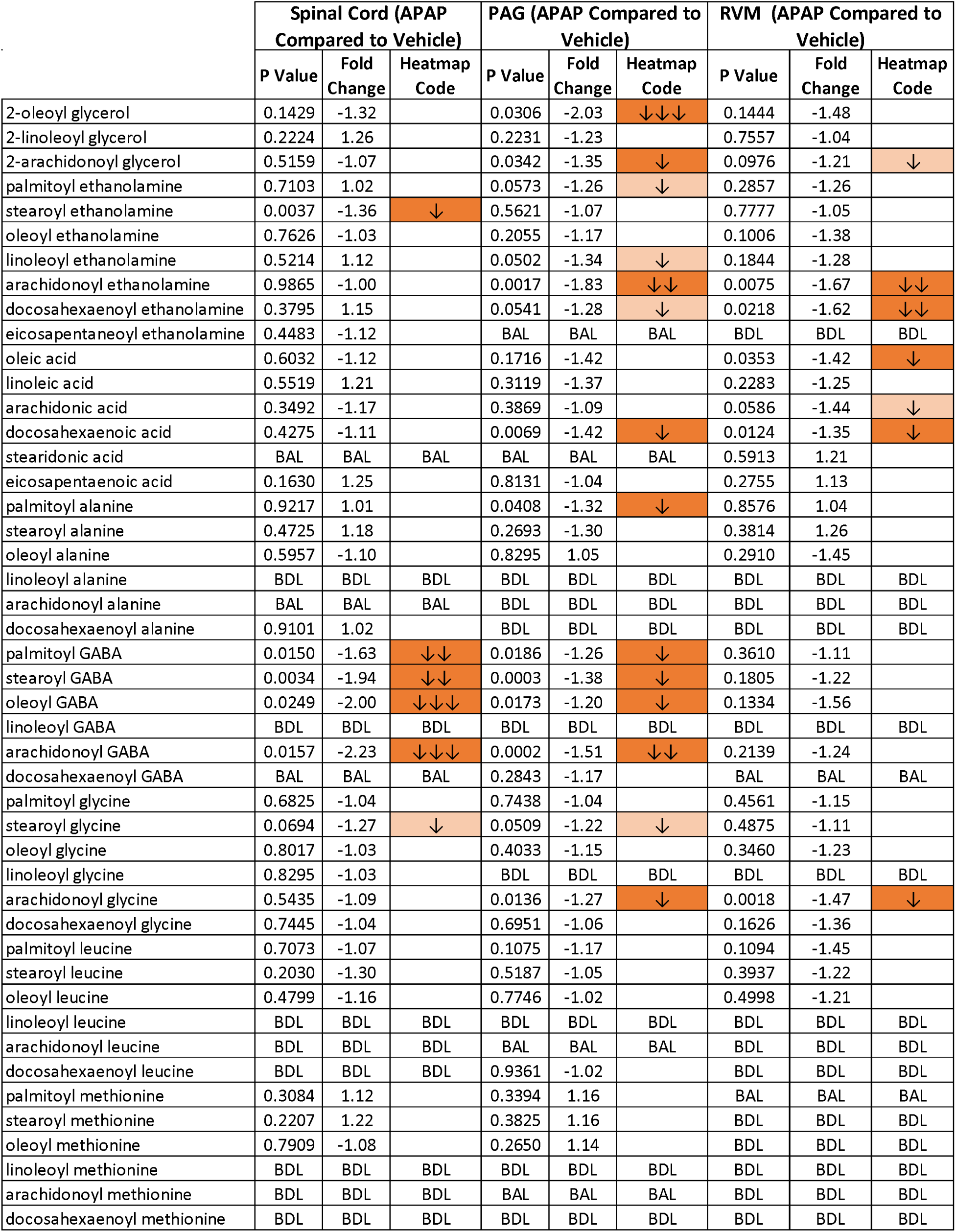
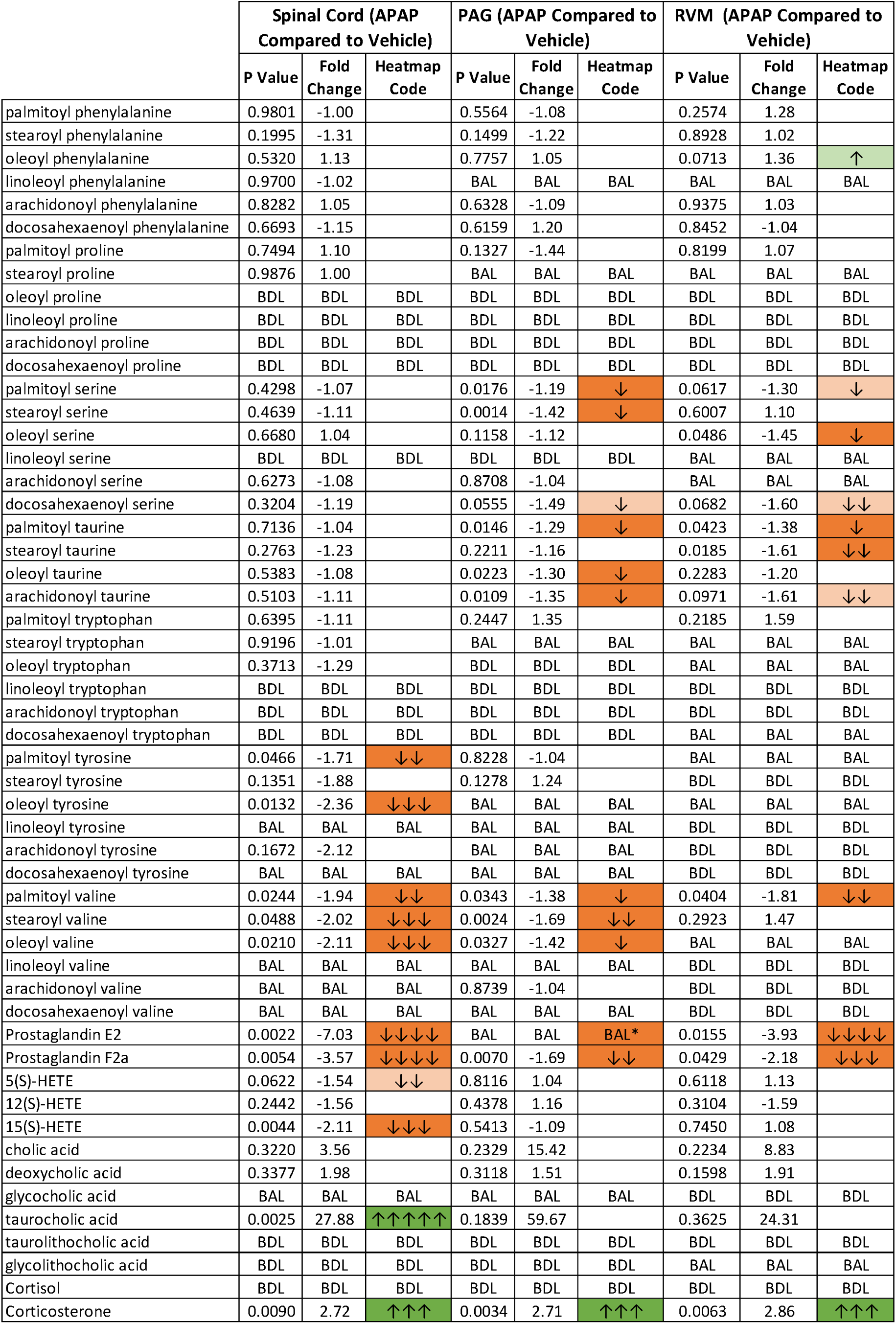
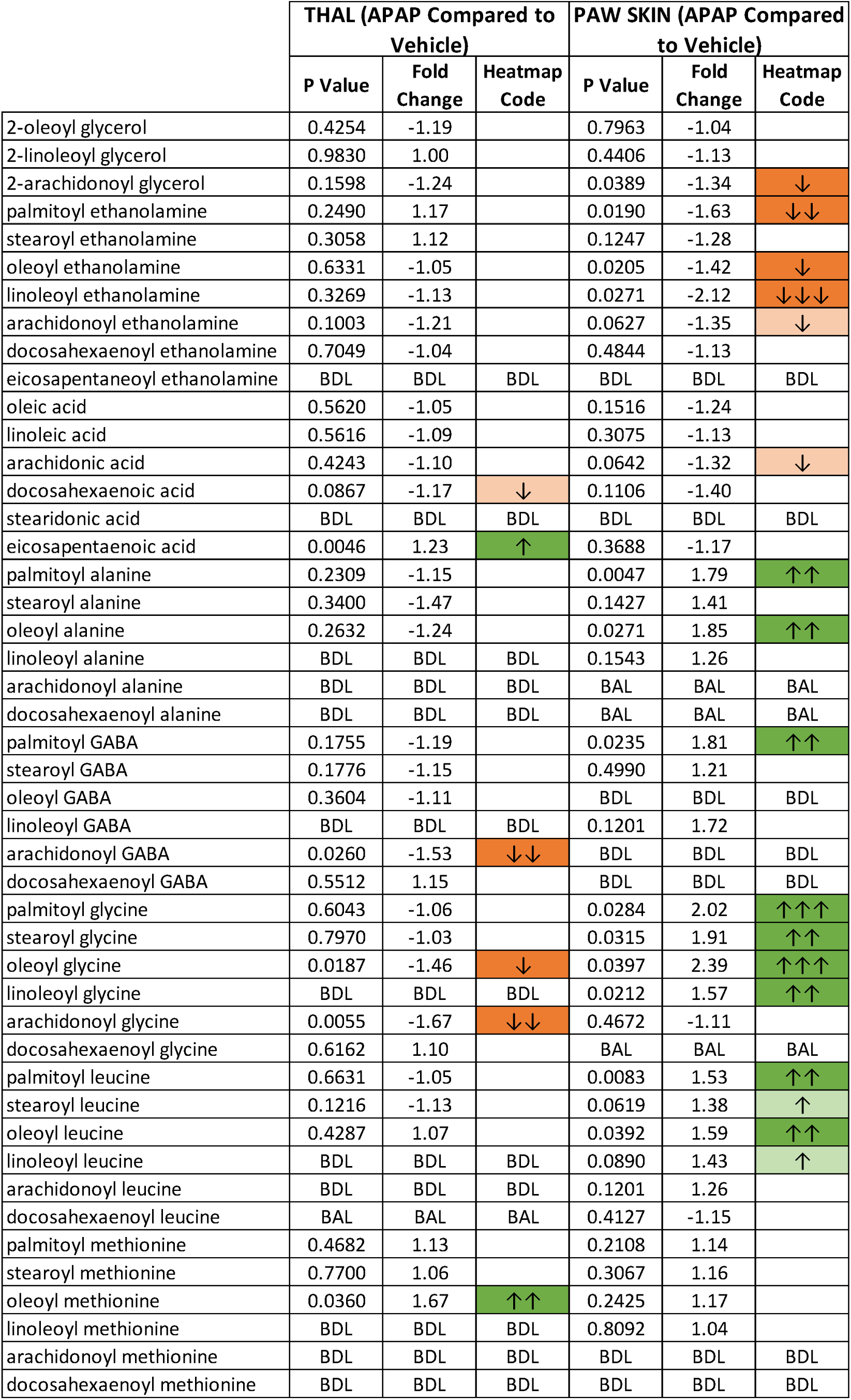
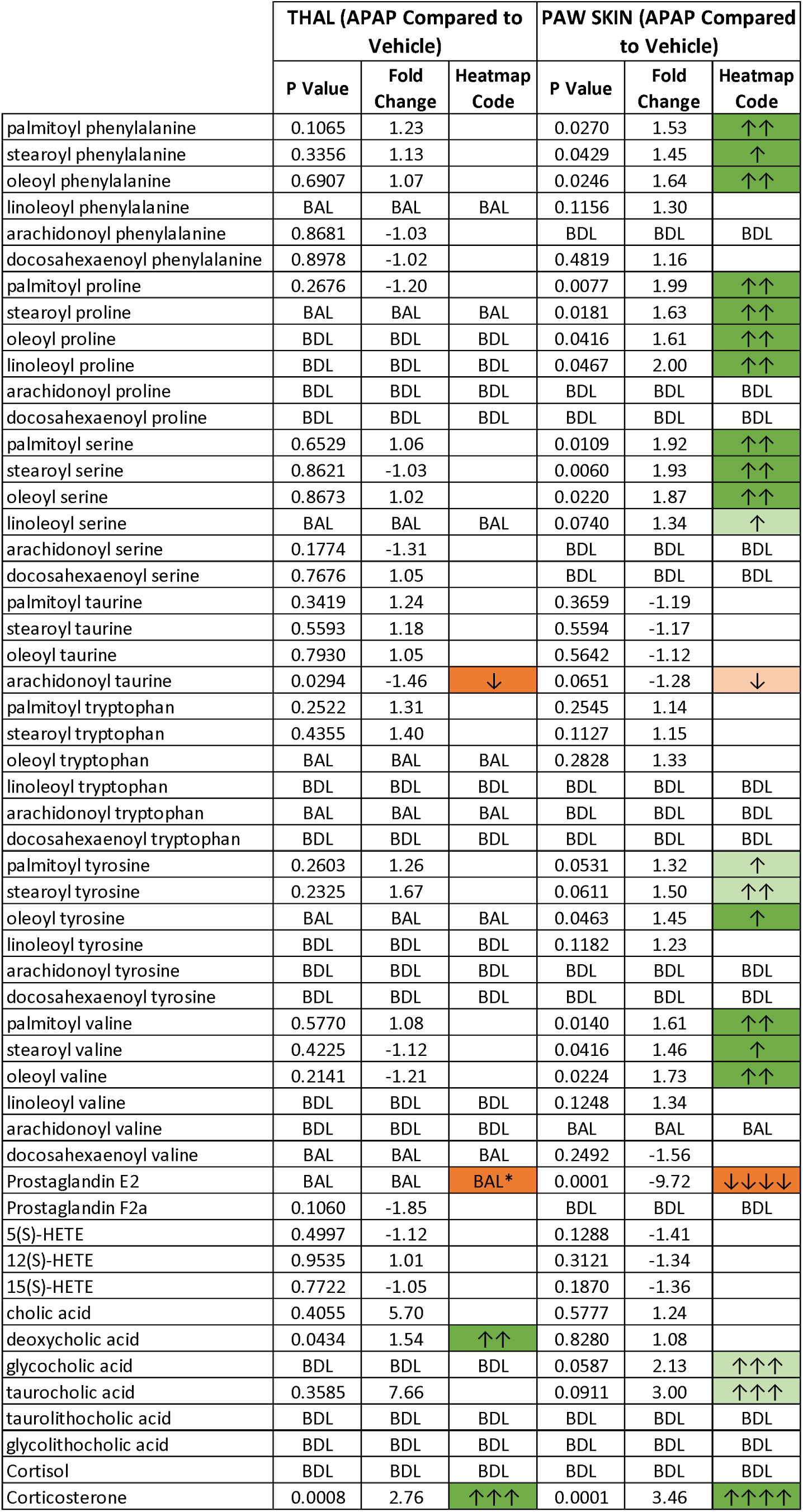
The full heatmap for all endogenous lipids detected at analytical levels in CNS and paw skin tissue. Dark green represents significant increases after treatment (p ≤0.05) while light green represents significance levels of p ≤ 0.1-0 >.05. Dark orange represents significant decreases after treatment (p ≤0.05) while light orange represents significance levels of p ≤ 0.1-0 >.05. Fold change is indicated by the number of arrows, where 1 arrow corresponds to 1–1.49-fold difference, 2 arrows to a 1.5–1.99-fold difference, 3 arrows to a 2–2.99-fold difference, 4 arrows a 3–9.99-fold difference, and 5 arrows a difference of tenfold or more. BAL (below analytical limits) refers to endogenous lipids that were measured in at least one sample but less than 4 making statistical analysis unreliable. BDL (below detectable limits) refers to endogenous lipids that were not detected in any samples. Blank cells indicate that endogenous lipids were present in at least 4 samples in each treatment group being analyzed and that no significant differences were measured.

