## Supplemental Material for "Acetaminophen alters endogenous lipid signaling and attenuates pathological pain through a mechanism requiring diacylglycerol lipase, monoacylglycerol lipase and cannabinoid CB1 receptors in mice"

**Supplemental Table 1**

| **Figure** | **Interaction** | **Treatment** | **Time** | **One-way ANOVA** |
| --- | --- | --- | --- | --- |
|  |  |  |  | **Antag./Inh effect at 30'** |
| 1A | F_6,45_=5.831, p=0.0001 | F _2,15_=8.477, p=0.0034 | F_2.391,35.87_=10.34, p=0.0001 |  |
| 1B | F_6,45_=0.8998, p=0.5034 | F_2,15_=0.0449, p=0.9561 | F_2.193,32.90_=0.7018, p=0.5155 |  |
| 1C | F_9,60_=4.293, p=0.0002 | F_3,20_=9.131, p=0.0005 | F_2.594,51.88_=1.678, p=0.1889 |  |
| 1D | F_9,60_=1.208, p=0.3074 | F_3,20_=35.91, p<0.0001 | F_2.614,52.28_=2.447, p=0.0819 |  |
| 2A | F_9,60_=3.210, p=0.0031 | F_3,20_=5.202, p=0.0081 | F_2.537,50.75_=6.549, p=0.0014 |  |
| 2B | F_9,60_=1.421, p=0.1995 | F_3,20_=10.07, p=0.0003 | F_2.189,43.77_=2.145, p=0.1250 |  |
| 2C | F_9,60_=3.387, p=0.0020 | F_3,20_=8.708, p=0.0007 | F_2.218,44.35_=6.206, p=0.0032 |  |
| 2D | F_9,60_=4.388, p=0.0002 | F_3,20_=8.442, p=0.0008 | F_2.066,41.33_=6.883, p=0.0024 |  |
| 2E | F_7.584,70.79_=5.151, p<0.0001 | F_3,28_=9.213, p=0.0002 | F_2.528,70.79_=8.649, p=0.0001 |  |
| 2F | F_8.892,82.99_=1.224, p=0.2924 | F_3,28_=0.4468, p=0.7215 | F_2.964,82.99_=0.7011, p=0.5524 |  |
| 3A | F_9,84_=11.23, p<0.0001 | F_3,28_=18.37, p<0.0001 | F_2.577,72.15_=1.879, p=0.1488 |  |
| 3B | F_9,84_=1.158, p=0.3322 | F_3,28_=0.6814, p=0.5708 | F_2.522,70.62_=1.048, p=0.3686 |  |
| 3C | F_3,27_=0.7625, p=0.5250 | F_3,27_=2.687, p=0.0664 | F_1,27_=24.34, p<0.0001 |  |
| 3D | F_3,27_=7.727, p=0.0007 | F_3,27_=4.366, p=0.0125 | F_1,27_=82.63, p<0.0001 |  |
| 3E | F_3,27_=6.037, p=0.0028 | F_3,27_=2.550, p=0.0767 | F_1,27_=75.76, p<0.0001 |  |
| 3F | F_3,27_=3.045, p=0.0459 | F_3,27_=0.0356, p=0.9908 | F_1,27_=40.01, p<0.0001 |  |
| 4A | F_9,75_=12.92, p<0.0001 | F_3,25_=9.418, p=0.0002 | F_2.112,52.81_=31.79, p=<0.0001 |  |
| 4B | F_9,78_=0.9100, p=0.5211 | F_3,26_=1.268, p=0.3060 | F_2.367,61.55_=1.553, p=0.2166 |  |
| 4C | F_3,25_=3.858, p=0.0214 | F_3,25_=1.277, p=0.3038 | F_1,25_=25.02, p<0.0001 |  |
| 4D | F_3,25_=19.87, p<0.0001 | F_3,25_=5.032, p=0.0073 | F_1,25_=152.7, p<0.0001 |  |
| 4E | F_3,25_=8.639, p=0.0004 | F_3,25_=5.571, p=0.0042 | F_1,25_=81.61, p<0.0001 |  |
| 4F | F_3,25_=5.062, p=0.0071 | F_3,25_=2.559, p=0.0777 | F_1,25_=44.09, p<0.0001 |  |
| 5A | F_9,66_=3.474, p=0.0015 | F_3,22_=16.43, p<0.0001 | F_2.154,47.38_=18.18, p<0.0001 |  |
| 5B | F_9,48_=2.522, p=0.0187 | F_3,16_=3.384, p=0.0442 | F_2.856,45.69_=15.10, p<0.0001 |  |
| 5C | F_9,66_=1.338, p=0.2347 | F_3,22_=0.3116, p=0.8168 | F_2.386,52.49_=2.840, p=0.0583 |  |
| 5D | F_9,84_=7.180, p<0.0001 | F_3,28_=14.42, p<0.0001 | F_2.811,78.81_=5.464, p=0.0023 | F_3,28_=1.300, p=0.2939 |
| 5E | F_9,81_=6.107, p<0.0001 | F_3,27_=12.40, p<0.0001 | F_2.642,71.33_=9.900, p<0.0001 | F_3,27_=1.774, p=0.1759 |
| 5F | F_9,81_=4.850, p<0.0001 | F_3,27_=14.98, p<0.0001 | F_2.466,66.59_=14.66, p<0.0001 |  |
| 5G | F_9,60_=3.641, p=0.0011 | F_3,20_=5.833, p=0.0049 | F_2.582,51.64_=9.367, p=0.0001 | F_3,20_=0.0125, p=0.9980 |
| 5H | F_9,78_=3.538, p=0.0010 | F_3,26_=12.39, p<0.0001 | F_2.314,60.16_=14.64, p<0.0001 |  |
| 6A | F_9,72_=4.183, p=0.0002 | F_3,24_=12.27, p<0.0001 | F_2.416,57.99_=9.10, p=0.0002 | F_3,24_=2.317, p=0.1011 |
| 6B | F_9,48_=3.813, p=0.0011 | F_3,16_=19.49, p<0.0001 | F_2.319,37.10_=3.315, p=0.0408 |  |
| 6C | F_9,60_=1.887, p=0.0713 | F_3,20_=9.825, p=0.0003 | F_2.857,57.13_=1.940, p=0.1360 | F_3,20_=2.106, p=0.1315 |
| 6D | F_9,57_=1.564, p=0.1485 | F_3,19_=11.39, p=0.0002 | F_2.523,47.94_=2.308, p=0.0982 |  |
| 6E | F_7.667,43.45_=1.681, p=0.1333 | F_3,17_=8.080, p=0.0015 | F_2.556,43.45_=2.769, p=0.0612 | F_3,17_=0.5968, p=0.6257 |
| 6F | F_6.430,42.87_=4.229, p=0.0016 | F_3,20_=7.967, p=0.0011 | F_2.143,42.87_=7.253, p=0.0016 |  |
| Sup.1A | F_9,60_=1.288, p=0.2626 | F_3,20_=1.342, p=0.2890 | F_2.626,52.52_=0.8230, p=0.4733 |  |
| Sup.1B | F_9,60_=0.3178, p=0.9661 | F_3,20_=20.79, p=0.3736 | F_2.524,50.49_=2.417, p=0.0869 |  |
| Sup. 2A | F_9,60_=1.189, p=0.3189 | F_3,20_=0.1950, p=0.8985 | F_2.851,57.02_=0.9214, p=0.4324 |  |
| Sup. 2B | F_9,60_=0.2853, p=0.9763 | F_3,20_=1.272, p=0.3108 | F_2.430,48.60_=2.609, p=0.0735 |  |
| Sup. 2C | F_9,60_=1.633, p=0.1263 | F_3,20_=0.1931, p=0.8998 | F_2.446,48.93_=1.354, p=0.2686 |  |
| Sup. 2D | F_9,60_=1.710, p=0.1064 | F_3,20_=0.7376, p=0.5419 | F_2.936,58.73_=0.8237, p=0.4839 |  |
| Sup. 3A | F_3,56_=0.5643, p=0.6408 | F_1,56_=2.295, p=0.1354 | F_3,56_=2.006, p=0.1236 |  |
| Sup. 3B | F_3,42_=0.4211, p=0.7388 | F_1,14_=0.1524, p=0.7022 | F_1.977,2768_=0.1558, p=0.8542 |  |
| Sup. 3C | F_1,14_=0.0063, p=0.9376 | F_1,14_=0.5245, p=0.4808 | F_1,14_=0.0570, p=0.8146 |  |
| Sup. 3D | F_1,14_=1.987, p=0.1804 | F_1,14_=0.0009, p=0.9761 | F_1,14_=5.463, p=0.0348 |  |
| Sup. 3E | F_1,14_=3.928, p=0.0675 | F_1,14_=0.6794, p=0.4236 | F_1,14_=5.591, p=0.0330 |  |
| Sup. 3F | F_1,14_=0.0550, p=0.8179 | F_1,14_=0.0002, p=0.9877 | F_1,14_=5.493, p=0.0344 |  |
| Sup. 4A | F_9,66_=0.4997, p=0.8695 | F_3,22_=0.7138, p=0.5542 | F_2.843,65.56_=0.9434, p=0.4213 |  |
| Sup. 4B | F_9,48_=1.025, p=0.4344 | F_3,16_=0.5173, p=0.6763 | F_2.576,41.22_=2.455, p=0.0850 |  |
| Sup. 4C | F_9,60_=1.673, p=0.1157 | F_3,20_=2.441, p=0.0942 | F_2.153,43.07_=2.578, p=0.0838 |  |
| Sup. 4D | F_9,84_=1.349, p=0.2247 | F_3,28_=0.1319, p=0.9403 | F_2.685,75.19_=1.310, p=0.2779 | F_3,28_=1.327, p=0.2856 |
| Sup. 4E | F_9,84_=1.203, p=0.3045 | F_3,28_=0.7344, p=0.5403 | F_2.947,82.51_=1.279, p=0.2871 | F_3,28_=0.3904, p=0.7608 |
| Sup. 4F | F_9,75_=0.3695, p=0.9461 | F_3,25_=2.164, p=0.1175 | F_2.915,72.88_=0.1801, p=0.9050 |  |
| Sup. 4G | F_9,60_=0.9353, p=0.5018 | F_3,20_=1.939, p=0.1557 | F_2.636,52.72_=0.3209, p=0.7846 | F_3,20_=1.069, p=0.3844 |
| Sup. 4H | F_9,78_=0.7765, p=0.6384 | F_3,26_=1.290, p=0.2987 | F_2.782,72.32_=0.7108, p=0.5386 |  |
| Sup. 5A | F_9,72_=0.3182, p=0.9665 | F_3,24_=1.260, p=0.3105 | F_2.707,64.96_=0.5988, p=0.6014 | F_3,24_=1.108, p=0.3654 |
| Sup. 5B | F_9,48_=1.053, p=0.4137 | F_3,16_=1.351, p=0.2933 | F_2.645,42.32_=0.290, p=0.8528 |  |
| Sup. 5C | F_9,60_=0.9860, p=0.4610 | F_3,20_=1.148, p=0.3540 | F_2.594,51.89_=0.1358, p=0.9179 | F_3,20_=1.513, p=0.2417 |
| Sup. 5D | F_9,57_=1.121, p=0.3633 | F_3,19_=1.677, p=0.2057 | F_2.415,45.88_=0.4300, p=0.6904 |  |
| Sup. 5E | F_7.022,39.79_=1.166, p=0.3439 | F_3,17_=4.085, p=0.0235 | F_2.341,39.79_=0.2194, p=0.8364 | F_3,17_=1.642, p=0.2170 |
| Sup. 5F | F_8.627,57.51_=0.4927, p=0.8675 | F_3,20_=0.6836, p=0.5724 | F_2.876,57.51_=0.3349, p=0.7918 |  |

**Supplemental Figure 1**

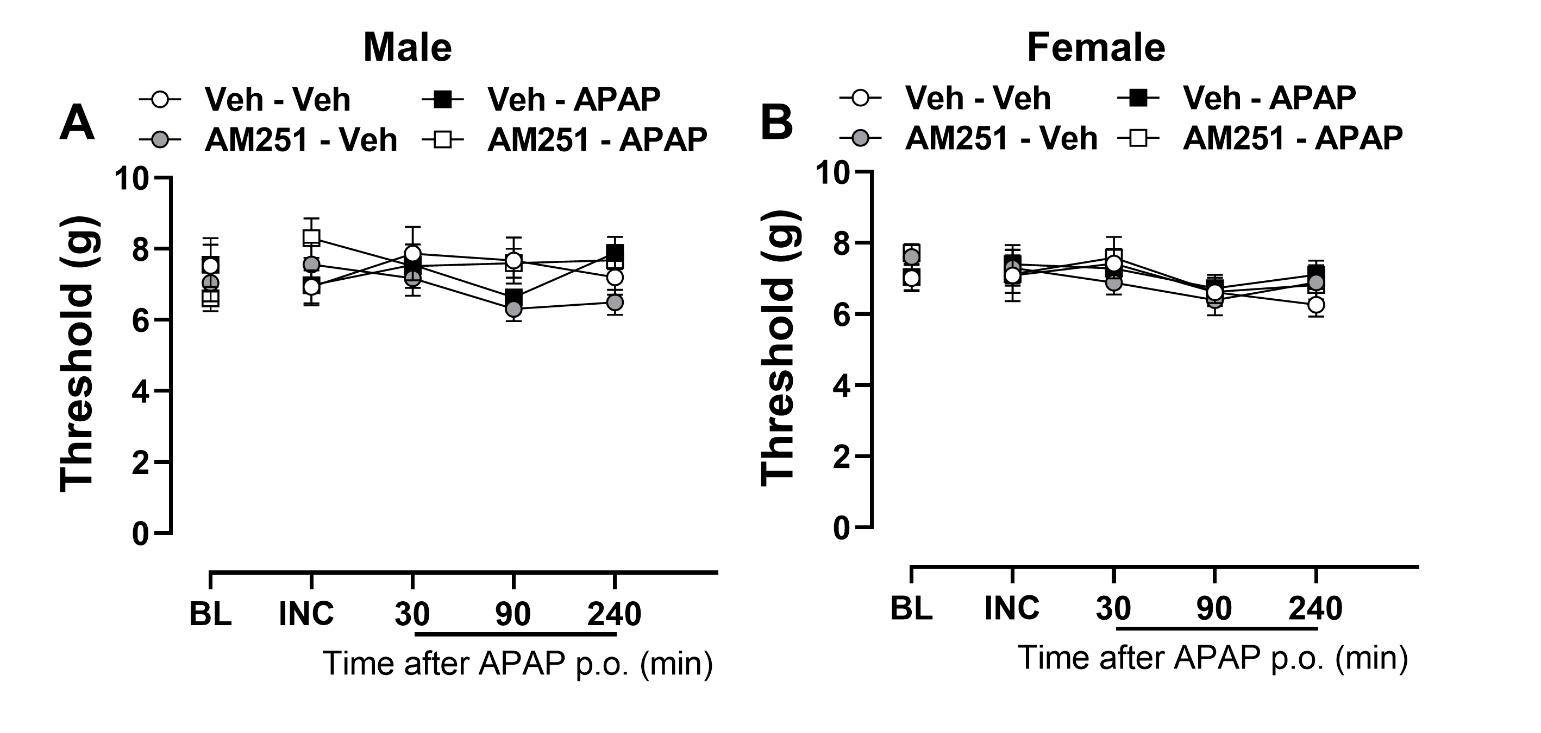

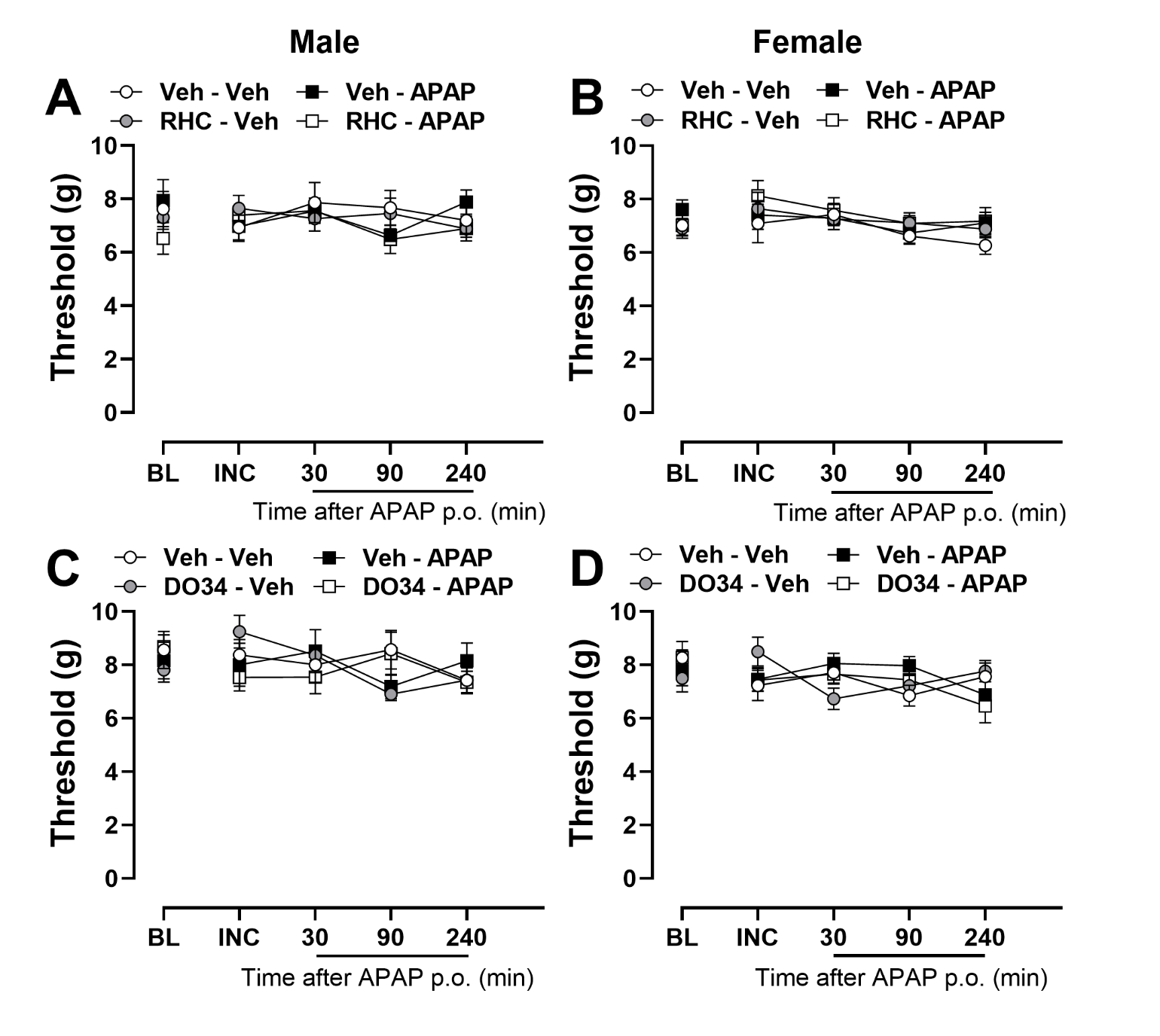
**Supplemental Figure 2**

**Supplemental Figure 3**

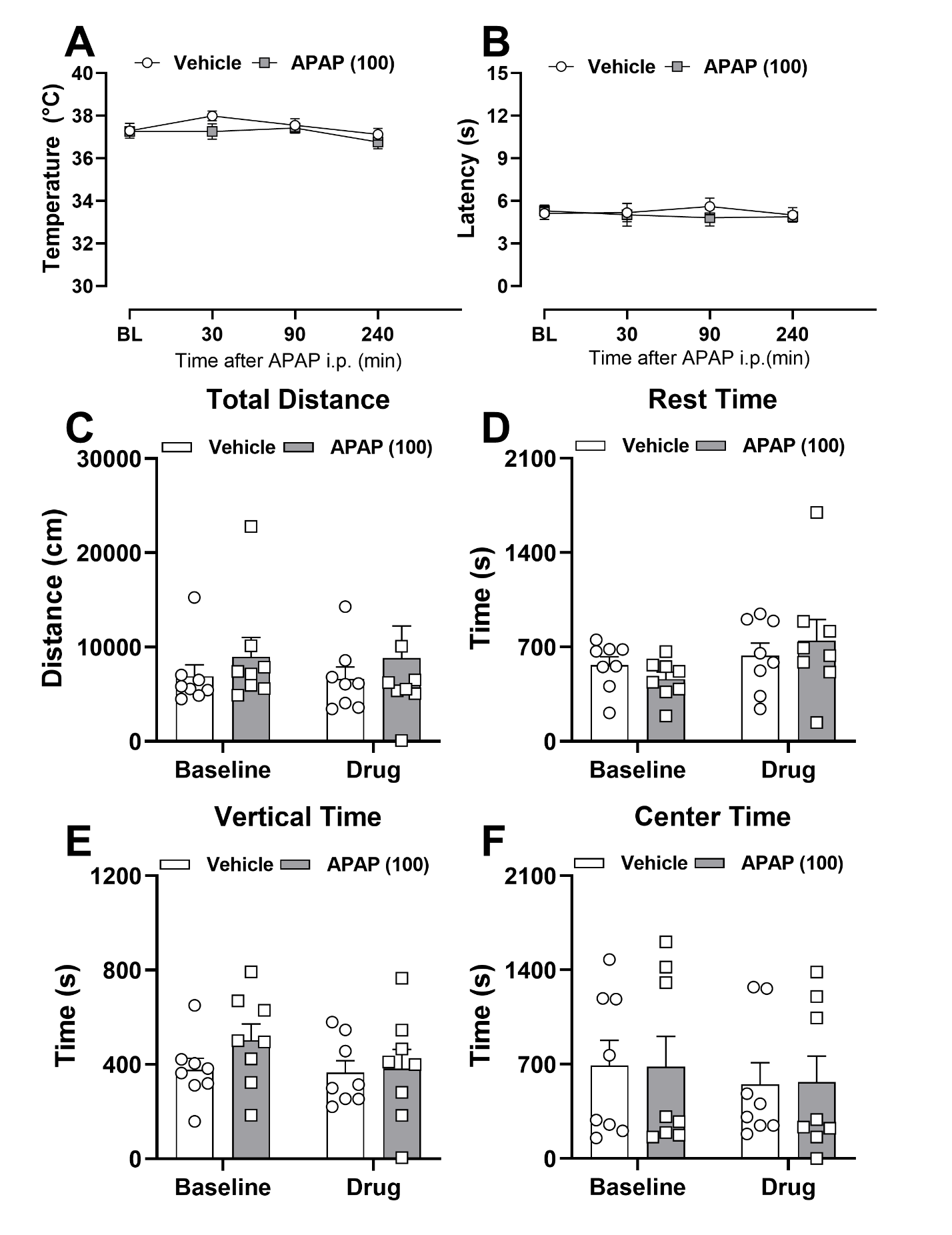

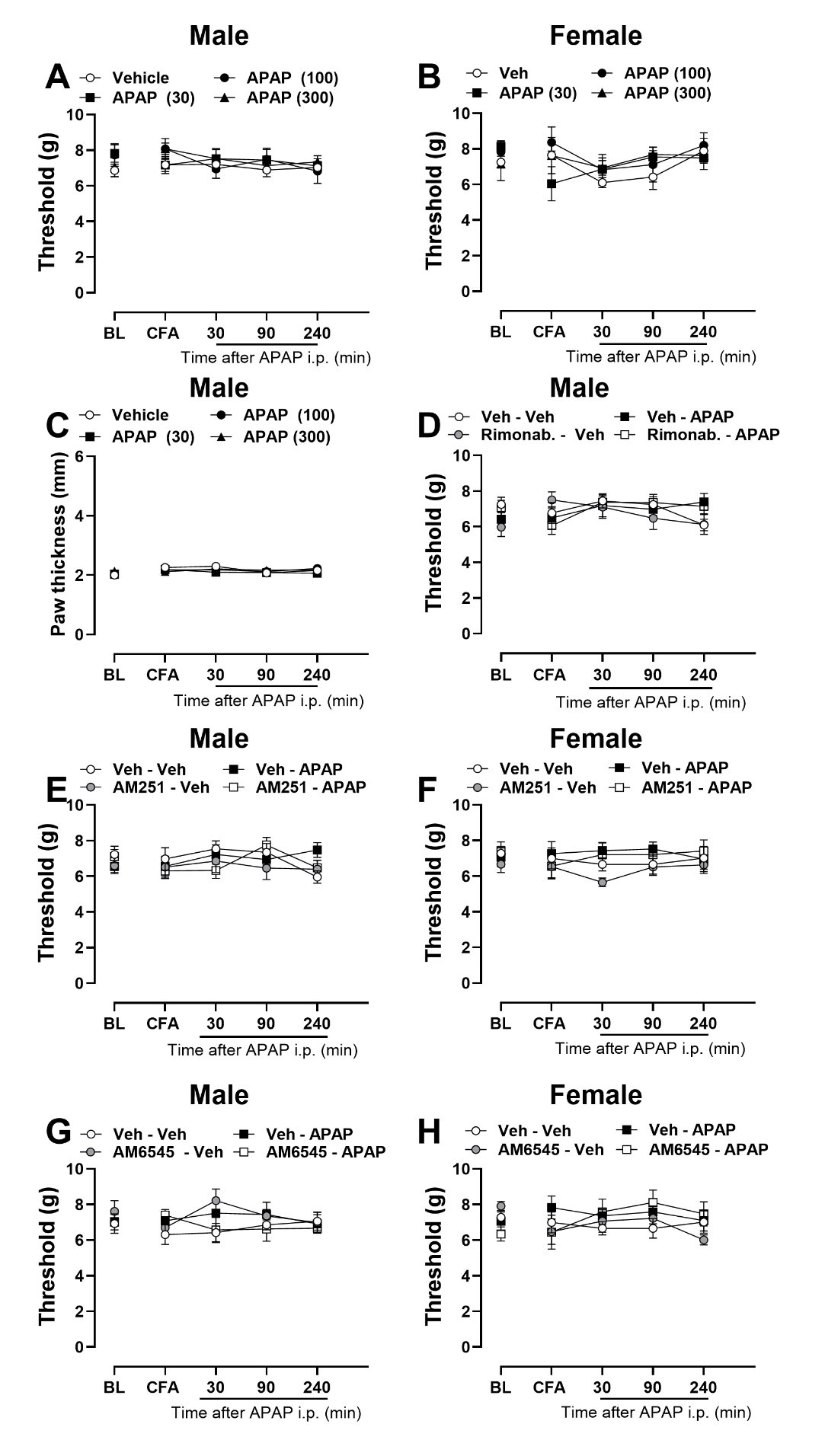
**Supplemental Figure 4**

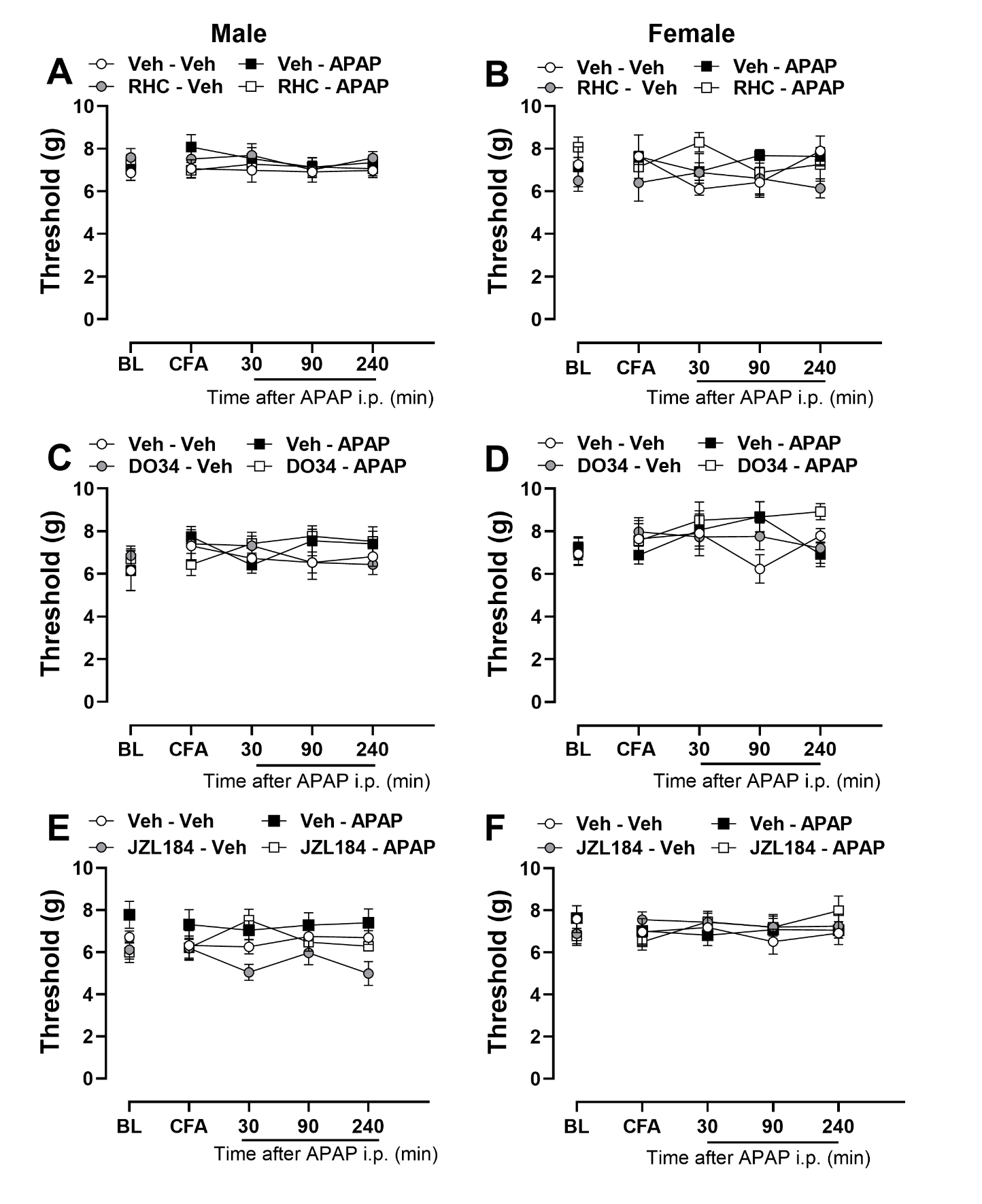
**Supplemental Figure 5**

**Supplemental Figure 6**

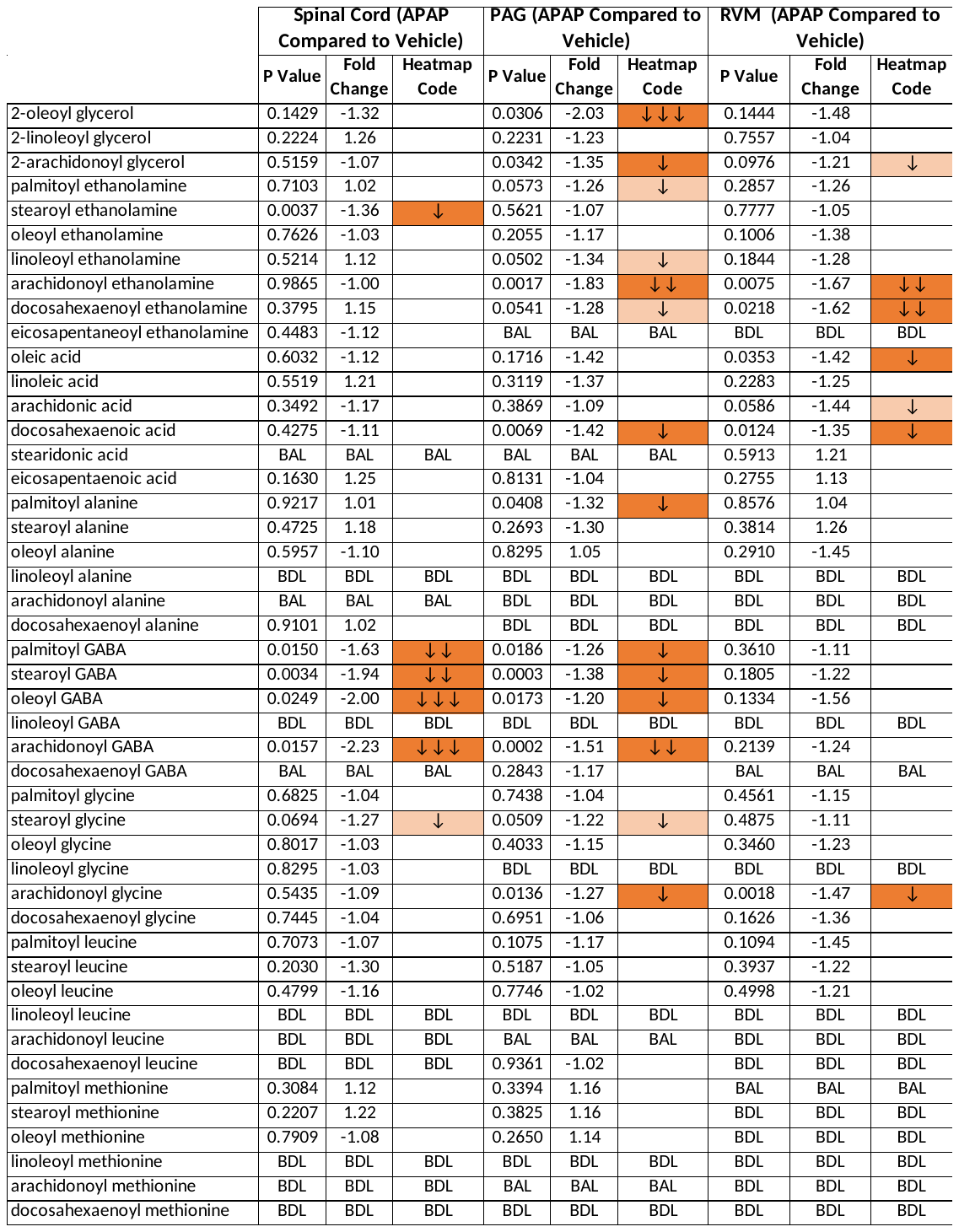

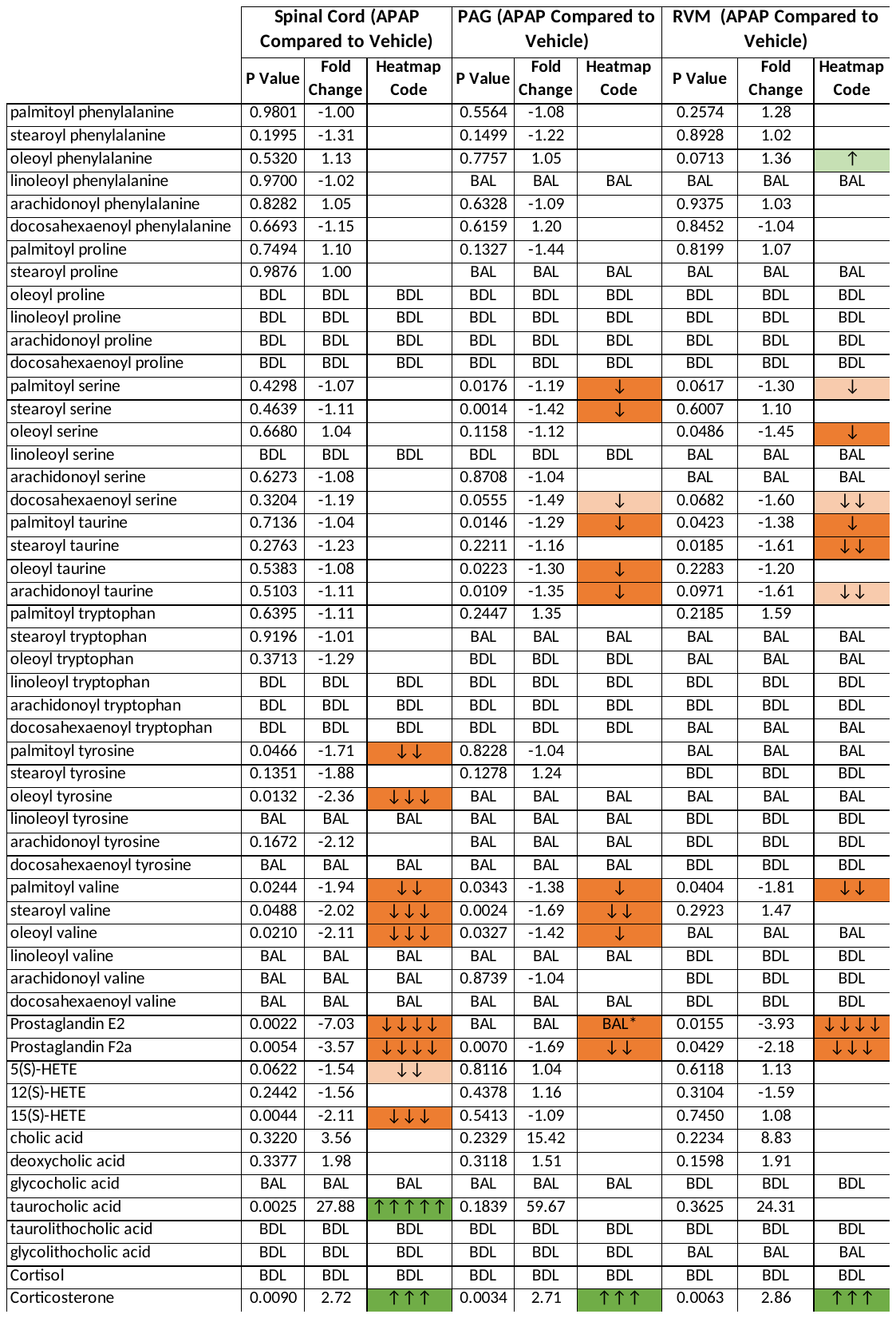

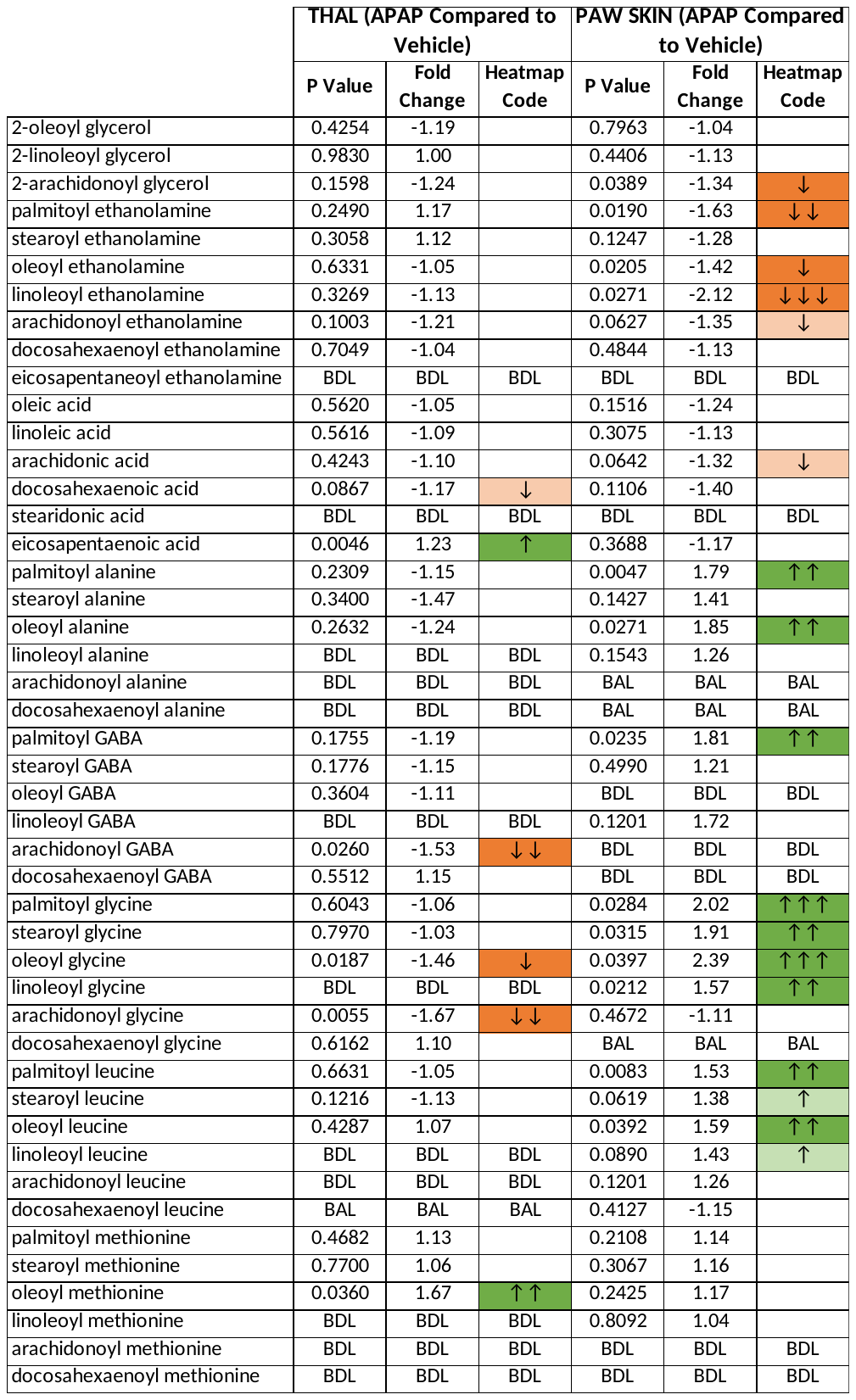

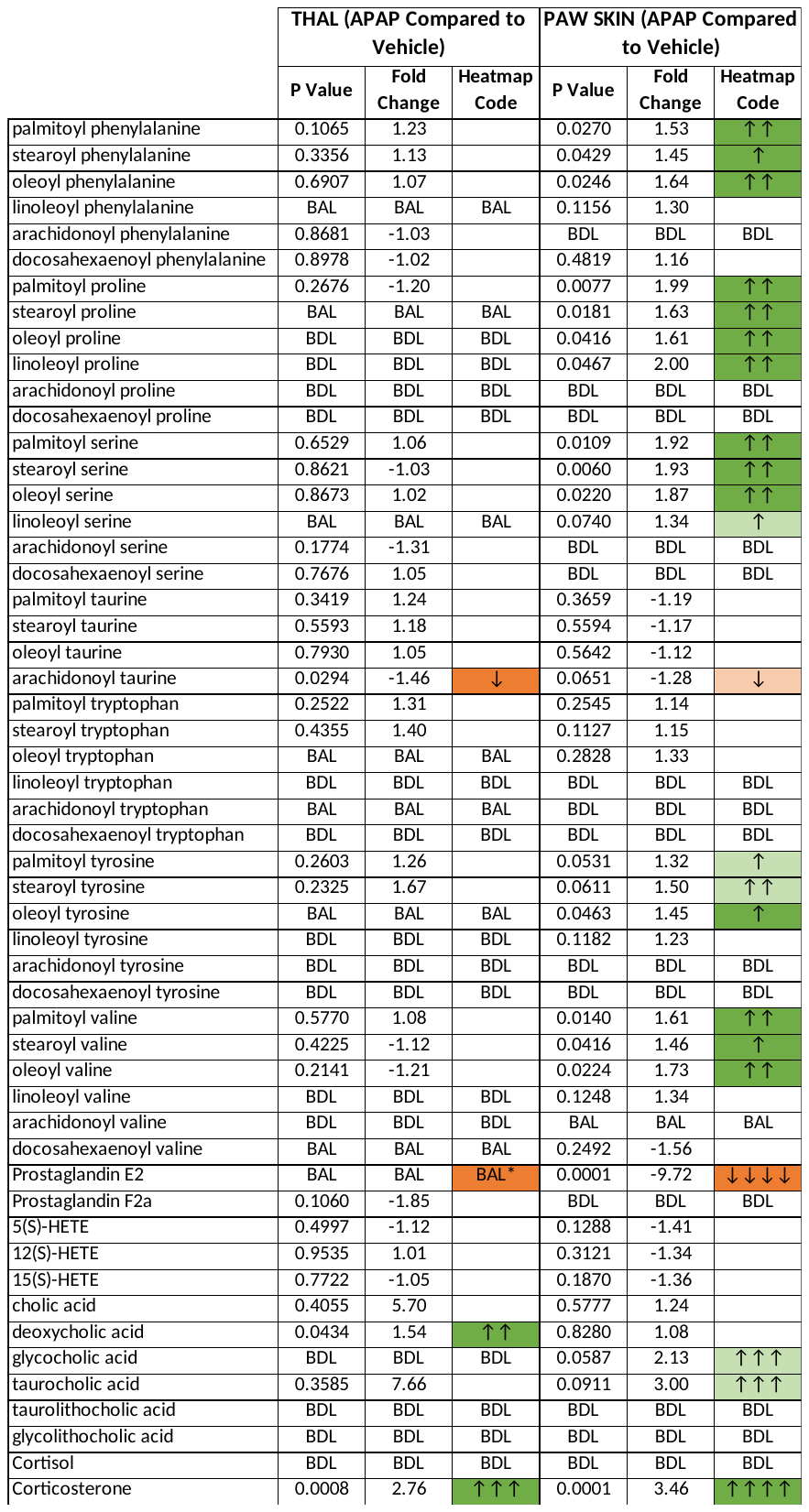
